# RosaSeed: Faster and Accurate Short Read Alignment Using a Configurable Seeding Strategy

**DOI:** 10.64898/2026.09.20.753062

**Authors:** Shyama M. Gandhi, Bruce F. Cockburn

## Abstract

Conventional short read DNA alignment algorithms, such as widely used BWA-MEM2, devote a substantial fraction of their execution time to seed generation. We present RosaSeed, a short read alignment algorithm, based on a configurable seeding framework, that significantly accelerates alignment while maintaining accuracy comparable to that of BWA-MEM2. RosaSeed replaces the BWA-MEM2 seeding kernel and directly supplies candidate seeds to the existing BWA-MEM2 chaining and alignment extension pipeline, producing standard SAM output suitable for downstream variant analysis. RosaSeed was evaluated on both simulated and real short read datasets, and its seeding time, fastqto-SAM time, alignment accuracy, and memory footprint were compared against BWA-MEM2, Minimap2, Bowtie2, and the Enumerated Radix Tree (ERT) algorithm. We also compared against ERT2, a proposed simplified variant of ERT that sacrifices a small amount of alignment accuracy to significantly increase the alignment throughput. With single-threaded software execution, our recommended RosaSeed configuration achieves 13.06× and 3.85× speedups in seed processing and fastq-to-SAM time, respectively, over BWA-MEM2. Corresponding seed processing (and fastq-to-SAM) speedups are 3.96×(2.48×) over ERT, 1.2×(1.44×) over ERT2, 4.82×(3.31×) over Minimap2, and 33.24×(10.83×) over Bowtie2. RosaSeed requires only 49.61 GB of peak memory, approximately 25% less than ERT and ERT2, while maintaining alignment accuracy comparable to that of BWA-MEM2. Following the recent release of minibwa, a prefetch-optimized FM index aligner, we additionally benchmarked RosaSeed against it on an upgraded 24-core, 48-thread AMD Zen 3 workstation. Our speed-optimized miniRosaSeed configuration achieves a 2.12× fastq-to-SAM speedup over minibwa for single-threaded execution while producing 2.58% higher standard accuracy. Against Strobealign, miniRosaSeed reported a 2.18× fastq-to-SAM speedup and 3.1% higher standard accuracy. For multi-threaded execution, miniRosaSeed was both faster and more accurate than minibwa for up to 26 software threads on the Zen 3 workstation. RosaSeed provides multiple runtime-memory configurations, spanning memory-efficient and high-performance strategies, enabling users to select operating points appropriate for diverse computational environments while maintaining alignment quality comparable to that of BWA-MEM2.

**The RosaSeed source code is publicly available at** https://github.com/smgandhi-18/RosaSeed.

## 1. Introduction

Over the past 30 years, dramatic reductions in the cost of deoxyribonucleic acid (DNA) sequencing, combined with advances in bioinformatics algorithms, have transformed biological and medical research. Genome reconstruction is required for *functional genomics*, when the location, function and expression of genes are determined [1]. The demand for human genome reconstruction has been driven by the creation of large genomic databases for medical research and public health management [2]. For example, in February 2024 the “All of Us” program, sponsored by the U.S. National Institutes of Health, announced that it had collected and made available to researchers a database containing over 245,388 diverse sequenced human genomes [3]. The United Kingdom hosts several large genome databases including the UK Biobank, which contains the genome sequences for almost 500,000 people [4]. Growing interest and successes in genome-based personalized medicine and consumer-driven genetic genealogical research are also adding to the demand for fast and inexpensive genome reconstruction. The release of more accurate reference genomes, such as the Telomere-to-Telomere (T2T) complete human genome assembly [5], as well as the increasing availability of references that are specific to sub-populations, are also growing the demand because of the additional genetic information that can be obtained by reprocessing existing sequenced genetic samples using more accurate and/or specific references.

On-going increases in short read sequencing throughput, together with falling sequencing costs and the growing demand for faster genome reconstruction, are straining available computational resources, making short read alignment and genome reconstruction a bottleneck in modern genomics pipelines [6, 7]. Contemporary short read alignment algorithms typically follow a multi-phase seed-and-extend pipeline consisting of (i) *seeding*, which identifies exact matches between segments in sequenced short reads and corresponding regions of the reference; (ii) *seed chaining*, which groups seeds with nearby aligning regions in the reference into longer candidate alignment regions; and (iii) *local alignment*, commonly implemented using the Banded Smith–Waterman (BSW) algorithm or other dynamic programming techniques [7, 8, 9]. Local alignment postulates likely mutations that allow seeds to be joined together to align the entire short read against the most likely corresponding positions in the reference genome. Among these stages, seeding plays a critical role: it determines both the quality of the candidate matches passed to the downstream stages and the processing latency through the pipeline. In BWA-MEM2, seeding alone accounts for approximately 40% of the total alignment runtime [10].

FM index based aligners, such as BWA-MEM [7], BWA-MEM2 [11] and Bowtie2 [6], perform backward search over a Burrows-Wheeler transformed reference [12]. Although this strategy enables compact indexing and a relatively small memory footprint, it requires base-by-base traversal of each short read using the FM index. The resulting irregular, and hence cache-resistant, memory access patterns into the large, precomputed FM index introduce a latency bottleneck, especially given the page-oriented architecture of modern high-density memory systems.

The ERT alignment algorithm [10] reduces this latency by replacing FM index traversal with a relatively large pre-computed radix tree data structure that enables multi-base lookups. Subramaniyan et al. [10] reported up to a 2.2× increase in seeding throughput at the cost of a substantially larger memory footprint (∼66.3 GB for the human genome) when compared to conventional software implementations of BWA-MEM2 (∼16.2 GB using a suffix array that is compressed by a factor of 8).

Minimizer-based aligners, such as Minimap2 [8], reduce the seeding overhead by indexing only a subset of representative *k*-mers (the minimizers) rather than all possible *k*-mers from the reference and short read sequences. This compressed sequence representation substantially reduces the seeding cost and has proven to be effective for long read alignment. However, because only a subset of candidate seeds is considered, minimizer-based approaches may miss some informative matches, resulting in reduced mapping accuracy.

A related subsampling strategy replaces exact *k*-mer seeds with longer, mutation-tolerant ones. Strobealign [13] subsamples the reference into *syncmers* [14] and links nearby pairs into gapped *strobemer* [15] seeds, attaining the uniqueness of much longer exact matches while remaining robust to sequence divergence. Unlike FM index aligners, it retrieves candidates by hash-based lookups into a precomputed seed table rather than using backward search. Using a new pipeline design, Strobealign employs its own chaining and extension strategy and does not integrate with an established downstream pipeline such as that of BWA-MEM2.

More recently, minibwa [16] pairs BWA-MEM style variable-length seeding with minimap2 chaining and SIMD alignment, hiding FM index cache-miss latency through timely software prefetching and batched short reads rather than reducing the number of memory accesses. It produces roughly twice the alignment speed of BWAMEM2 with comparable accuracy, while further reducing downstream cost by capping the number of chains per read and reducing the time spent aligning reads to highly repetitive regions of the reference.

Despite these advances, the existing algorithms typically improve alignment speed by sacrificing memory efficiency and/or seeding accuracy. Consequently, an opportunity remains for methods that substantially speed up FM index traversal without incurring a large memory footprint, while preserving both alignment quality and compatibility with existing short read alignment pipelines.

In this work, we present *RosaSeed*, a fast and accurate short read alignment algorithm that is based on a configurable seeding method. The name honors the contribution of Dr. Rosalind Franklin to the discovery of the structure of DNA. RosaSeed accelerates seed generation using an *s*-base reference genome (where *s* ≥ 1), a compressed occurrence array, and a *k*-mer jump table that finds all alignment positions for a short read segment that contains exactly *k* nucleotides. The seeding method supports multiple symbol granularities (that is, *s* values) in which either one, two or three DNA bases are encoded per FM index symbol. RosaSeed further introduces multiple supplementary seeding strategies, including fixed-pivot seeding and adaptive gap-directed seeding, to improve the coverage of the generated seeds (and hence increase the alignment accuracy) while still producing fast seed generation. The resulting seeds are transformed back to base-level reference positions and forwarded to the proven BWAMEM2 chaining and alignment-extension pipeline, while providing significantly reduced seed generation time. By accelerating the seed generation stage without replacing the downstream stages of the alignment pipeline, RosaSeed provides a drop-in alternative to the conventional BWA-MEM2 seeding stage while exposing configurable memory-performance trade-offs through its multiple adjustable algorithm parameters and heuristic thresholds.

The main contributions of this work are as follows:

- With RosaSeed we introduce an *s*-base FM index that is computed from a reference that is derived from the concatenation of *s* ≥ 2 *s*- base references that are phase-shifted with respect to each other. Including the multiple phase-shifted *s*-base references ensures that all possible seed boundaries are considered. The proposed seeding framework also incorporates an initial jump table look-up that speeds up the search for seeds.
- We propose an initial strand-aware disjoint pivot sweep (Phase A) that identifies the longest possible seeds while avoiding overlap between successive seeds. This is achieved through adaptive *s*- base index extension and reference walking (comparing the short read with a suspected alignment location in the reference) in a way that efficiently explores candidate seed intervals while preserving strand consistency.
- We propose configurable supplementary seeding strategies that complement the primary sweep through either fixed-pivot seeding (Phase B) or adaptive gap-directed seeding (Phase C), improving seed coverage in the read not covered by Phase A while providing flexible trade-offs between runtime and seed density.
- We incorporate configurable suffix array compression into RosaSeed, optimizing an important memory-performance trade-off through sampled suffix array compression.
- We integrate RosaSeed with the existing BWA-MEM2 chaining and alignment stages, preserving compatibility with proven seed chaining and local alignment methods. Optional pre-chain seed filtering heuristics are proposed that reduce the propagation of low-value seeds into chain construction to avoid slowing down the BSW stage while not significantly lowering alignment accuracy.
- We describe the results of a comprehensive experimental evaluation on simulated and real human short read datasets using the complete T2T-CHM13v2.0 reference genome, comparing multi-threaded implementations of RosaSeed against BWAMEM2, ERT, ERT2, Bowtie2 and Minimap2 with respect to runtime, memory footprint, energy consumption, scalability, and multiple alignment accuracy metrics.
- We also propose our speed-optimized miniRosaSeed configuration of RosaSeed, benchmarked against minibwa and Strobealign on our newer 24-core AMD Zen 3 workstation. We also evaluate downstream variant-calling consistency using DeepVariant [17] and present the results.

The remainder of this article is organised as follows: Section 2 reviews background on FM index based alignment and related seeding algorithms. Section 3 describes the RosaSeed algorithm in detail, including the *s*-base derived reference, FM index construction, jump table design, and the multi-phase seeding stage comprising Phase A and followed by either Phase B or Phase C. Section 4 presents experimental results and comparisons with other major alignment algorithms. Section 5 discusses performance-cost trade-offs available with different RosaSeed configurations. Finally, Section 6 concludes the article and proposes directions for future work.

## 2. Background

### 2.1. Basic Terminologies

This section reviews the genomic and algorithmic concepts used throughout this article. Comprehensive surveys of short read alignment methods are provided in [18, 19].

The genetic blueprint of an organism is its *genome*, which (apart from some viruses) is encoded in long DNA polymers composed of four nucleotide types that differ in their *base* functional group, which is one of: adenine (A), cytosine (C), guanine (G) and thymine (T). One human genome consists of a sequence of approximately 3.2 billion nucleotide pairs. A DNA polymer has a twisted ladder structure, called a double helix, where each ladder rung is constrained by inter-base hydrogen bonds to be one of the four ordered base pairs AT, TA, CG and GC. The bases A and T are *complements* of each other, as are C and G. The two ladder rails have opposite chemically distinguishable directions so that the base sequence of the nucleotides along the forward (reverse) direction of one rail is the *reverse complement* of the base sequence along the forward (reverse) direction of the other rail.

*Next Generation Sequencing* (NGS) machines use massively parallel processing to rapidly and cost-effectively determine the sequence of approximately 150 bases at the two ends of hundreds of millions of DNA fragments, which are cleaved chemically at random positions from a purified DNA sample [20]. The recovered 150-base sequences are called *short reads. Short read alignment* is the key problem that must be solved 100s of millions of times to reconstruct an organism’s genome, given a reference genome for the species, for the corresponding large set of short reads produced by an NGS machine. Each short read must be aligned to the best-matching, and thus most accurate (as determined by standard genetic cost functions [9]) positions in the reference, allowing for possible structural mutations such as nucleotide substitutions, insertions, deletions, and more complex nucleotide rearrangements that distinguish the organism’s particular genome from the reference. Identifying these mutations is essential for understanding the genetic mechanisms underlying healthy and diseased cell function, developing personalized medical treatments, and detecting cancerous cells [20, 21] and extracellular DNA fragments [22].

### 2.2 The seed-and-extend alignment pipeline

Most contemporary short read alignment algorithms decompose the alignment problem into three successive stages, following the seedand-extend paradigm [7, 8] (Figure 1). In the first stage, seeding, the aligner identifies short exact matches between substrings of the read and corresponding substrings in the reference genome. These matching substrings, called *seeds*, serve as anchors that constrain where alignments of the full read are subsequently sought in the reference. In the second stage, *seed chaining*, groups of compatible seeds are identified and placed into sequences (that is, *chains*) such that the seeds appear in the same order as the corresponding regions in the reference. This ordering reduces the relatively expensive dynamic programming work that must subsequently be performed. In the third stage, *local alignment*, the seeds in each candidate region are precisely aligned against the reference using BSW dynamic programming [9], which considers possible nucleotide substitutions, insertions and deletions, producing a CIGAR-encoded alignment [23] that specifies the most likely alignment of each read. The final output is a SAM file [23] suitable for downstream variant calling using tools in standard genomics packages, such as GATK [24, 25].

**Figure 1.**
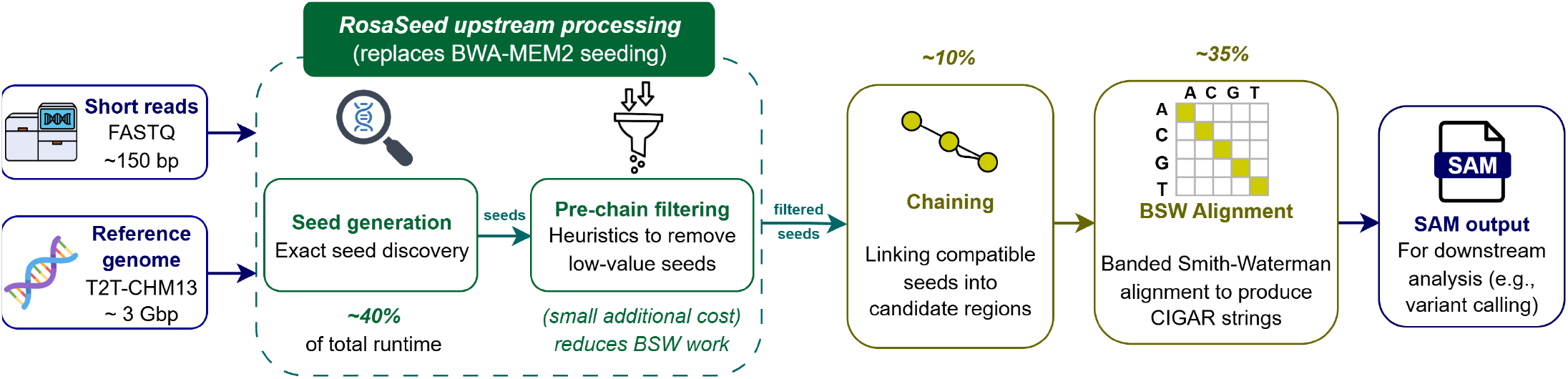
Integration of RosaSeed into the BWA-MEM2 workflow. RosaSeed replaces the original seed-generation stage followed by optional pre-chain seed filtering to reduce redundant and low-value seeds before chaining. The resulting seed set is passed to the standard BWA-MEM2 chaining and BSW alignment modules to generate read alignment records in the standard SAM file format. Runtime percentages are approximate and reflect the dominant runtime components of BWA-MEM2; the remaining execution time is attributable to output generation, index loading and auxiliary processing.

The seeding stage is typically the dominant computational bottleneck in the pipeline. In BWA-MEM2, seeding accounts for approximately 40% of the total runtime, chaining for around 10%, and BSW for around 35% [10]. This unevenness arises because seeding must process every base of every read against a compressed genome-wide index, generating millions of irregular (and hence relatively slow) memory accesses, while chaining and BSW operate only on the much smaller set of candidate seed sequences that are output from seeding. RosaSeed addresses this bottleneck by replacing BWA-MEM2’s seed generation stage with a carefully simplified and optimized new seed generation stage that includes lightweight pre-chaining heuristics, while preserving the established and trusted chaining and BSW alignment stages from BWA-MEM2 (Figure 1).

### 2.3. FM index and SMEM-based seeding

The FM index [12] is the core data structure underlying fast seeding in BWA-MEM, BWA-MEM2 and Bowtie2 that is constructed from the *Burrows-Wheeler Transform* (BWT) of the reference genome. The BWT is a reversible permutation of the reference sequence obtained after lexicographically sorting all of the cyclic rotations of the reference and then concatenating the last symbol of every rotation [12]. The *FM index* comprises three auxiliary data structures derived from a BWT: the *occurrence array* Occ(*σ, i*), which stores the cumulative number of occurrences of symbol *σ* in BWT[0 … *i*) (denotes the first *i* elements at positions *i* = 0, …, *i* − 1 of the BWT of the given reference); the *count array C*[*σ*], which stores the number of symbols in the reference genome lexicographically smaller than a given symbol *σ*; and the *suffix array* (SA), which contains a unique reference position for each BWT index value. Together, these data structures support fast *backward search* for strings, according to their suffix, in the reference: given a short read query string *R*, the FM index is searched by repeatedly updating two index values ℓ and *h*, where ℓ ≤ *h*, while scanning along *R* one base at a time, from right to left, along longer suffixes of the query. The low, ℓ, and high, *h*, index values are initialized to 0 and the highest index into the SA, respectively. At any point in the backward search, the index interval [ℓ, *h*) specifies the SA values SA[ℓ], …, SA[*h* − 1] that point to all the locations in the reference that are exact matches with the symbols scanned so far in *R*. Each backward search extension requires two queries into the large occurrence array, namely Occ(*σ*, ℓ) and Occ(*σ, h*), together with one lookup *C*[*σ*] [12]. Because the large occurrence array is accessed with addresses that do not follow a simply predictable order, these lookups exhibit poor cache locality and so lookups will produce relatively slow, last-level cache misses. For a 150 base pair (bp) read, BWA-MEM2’s FM index implementation requires approximately 68 KB of Occ(*σ, i*) data per read, where approximately 40% of accesses result in cache misses [10].

BWA-MEM2 generates seeds for each short read using the *Super-Maximal Exact Match* (SMEM) algorithm [7], illustrated in Figure 2. An exact match between a read substring and the reference is called a *Maximal Exact Match* (MEM) if it cannot be extended in either direction without introducing a mismatch. A MEM is called *supermaximal* if it is not fully contained within any other MEM. The SMEM algorithm proceeds in two passes. In the *forward search pass*, the aligner places a *pivot* (the current scanned position in the read) in the read and extends the match one base symbol at a time going to the right, at the same time recording each position where the BWT index interval changes size as a *Left Extension Point* (LEP). Each LEP marks a boundary where the hit set (the SA interval) changes, and the forward MEMs up to that point are stored for use in the subsequent backward pass. In the example of Figure 2, the pivot is placed at base T (shown in orange). Forward extension from the pivot produces the MEMs TA, TAC, TACC and TACCTCC, which are all stored for backward extension. In the *backward search pass*, the aligner extends each stored MEM to the left, one base at a time, until the SA [ℓ, *h*) interval either collapses to empty [ℓ, ℓ) or the leftmost end of the read is reached. MEMs that are fully contained within a longer match are discarded (marked with × in Figure 2), and the remaining non-contained matches are reported in a set of found SMEMs. In the figure, AGGTA and AGGTACC are shown discarded as they are contained, while ACGCTTAGGTAC and GCTTAGGTACCTCC are retained as the final SMEMs. The pivot is then advanced leftward to one base beyond the leftmost base of the longest SMEM just found, and the entire process is repeated until the read is fully scanned. This sweep is performed at multiple reseed thresholds to find seeds in repetitive DNA regions that the primary pass might have missed [7].

**Figure 2.**
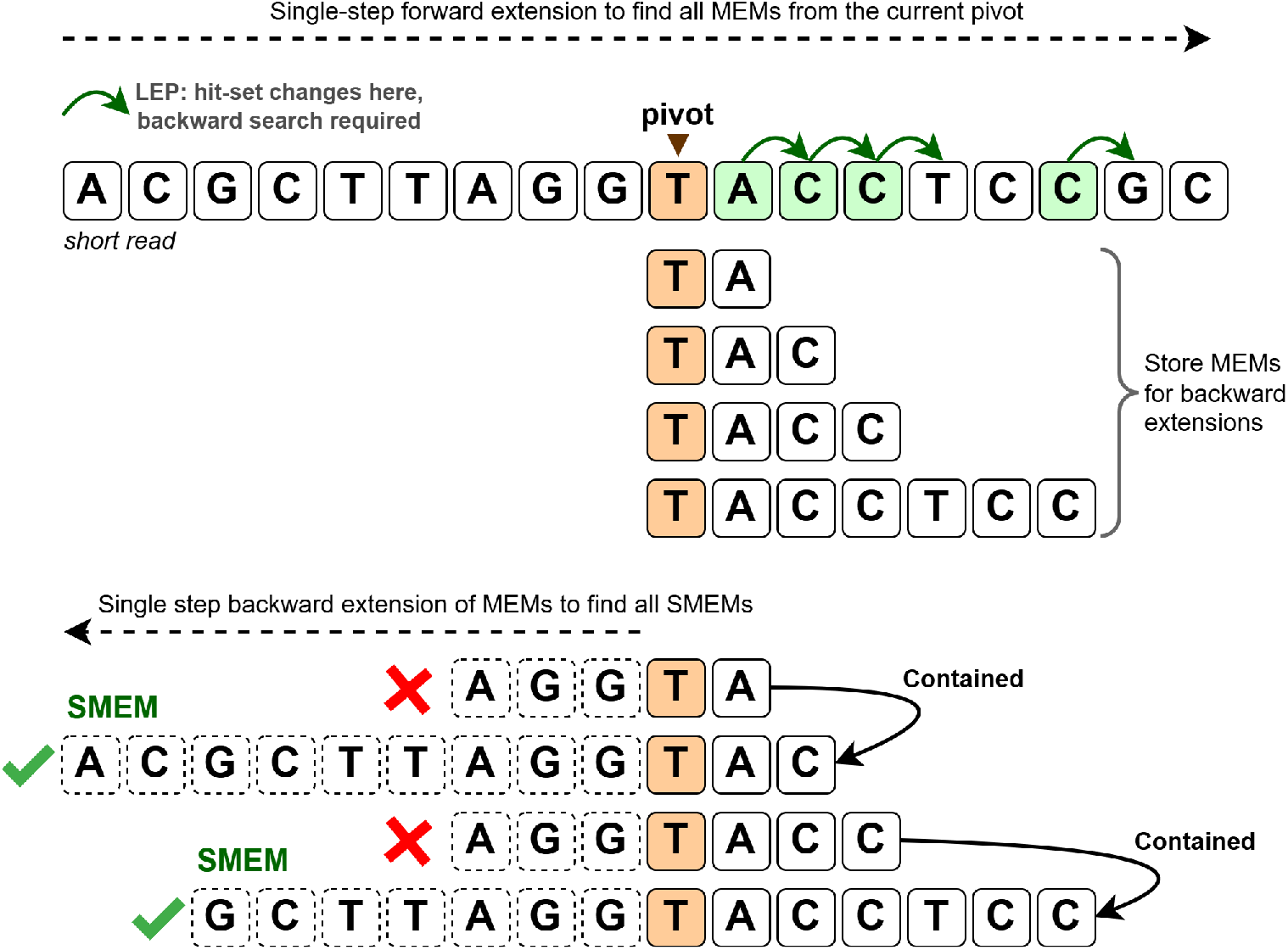
The SMEM seeding algorithm used by BWA-MEM2. Top: The forward search pass starts at the current pivot (orange) and extends going right one base at a time, storing MEMs at each Left Extension Point (LEP) for backward extension later on. Bottom: The backward search pass extends each stored MEM to the left. MEMs that are found to be fully contained within a longer match (×) are discarded, and the remaining non-contained matches are reported as final SMEMs (✓).

### 2.4. Related seeding approaches

As described below, several different approaches have been proposed to address the seeding bottleneck, each making different tradeoffs between speed, memory footprint and compatibility with the downstream stages of the standard alignment pipeline.

#### BWA-MEM (2013) and BWA-MEM2 (2019)

BWA-MEM [7], which introduced the SMEM-based seeding algorithm described in Section 2.3, appears to be the most widely employed short read aligner. It is also the standard aligner used in the GATK Best Practices pipeline [25]. BWA-MEM2 [11] accelerates BWA-MEM using SIMD-optimized FM index operations (where SIMD refers to the parallel *single instruction, multiple data* instruction available in many modern computers) and a reduced suffix array compression factor, achieving approximately 1.3–3.1× speedup over the original algorithm. BWA-MEM2’s default index uses approximately 16.2 GB of memory using a SA compression factor of 8. On the other hand, BWA-MEM2, when using an uncompressed suffix array, requires approximately 41.2 GB of memory and is used as the baseline short read aligner in this work. The corresponding seeding stage remains FM-index-based and assumes base-by-base alignment, and continues to account for approximately 40% of the runtime.

#### The Enumerated Radix Tree (ERT) Algorithm (2021)

ERT [10] addresses the data bandwidth bottleneck by replacing the FM index with a relatively large radix tree data structure, derived from the reference, that supports multi-base lookups. The precomputed ERT data structure enumerates all 4^15^ possible 15-mer prefixes of candidate seeds. Each lookup fetches the corresponding radix tree root in a single memory access, allowing *k* = 15 consecutive bases to be matched in exactly one operation. The CPU-ERT version of ERT achieves up to 2.2× higher seeding throughput than BWAMEM2 in software. The trade-off is a substantially larger index memory footprint: approximately 66.3 GB for a human genome, compared to 16.2 GB for BWA-MEM2. This relatively large footprint would be prohibitive in many contemporary workstations but would be a feasible option for cloud-based servers. A *Field-Programmable Gate Array* (FPGA) accelerated version of ERT achieves 3.3× seeding throughput over the software version of ERT.

#### Minimap2 (2018)

Minimap2 [8] generates seeds using *minimizers*: for each window of *w* consecutive *k*-mers, only the lexicographically smallest *k*-mer is retained as one representative minimizer for the window. This simple heuristic dramatically reduces the number of considered seeds and subsequent lookup operations, making Minimap2 especially efficient for long read alignment, where seed density is relatively low. For short reads, however, the minimizer approach introduces speed-performance trade-offs because important seeds at any given position may be omitted if a neighbouring *k*-mer happens to be smaller. In our experiments, Minimap2 was run using its short read preset configuration (`-ax sr -k19 -w10`), which is designed for short Illumina reads [8]. Despite this setting, Minimap2’s total alignment time is less than that of BWA-MEM2 as the minimizer heuristic inherently processes fewer seeds per read than the SMEM-based approach of BWA-MEM2.

#### Bowtie2 (2012)

Bowtie2 [6] generates seeds using a combination of FM index backward search and a heuristic seed interval strategy that places seeds at fixed intervals along the read. It is especially memory-efficient (4.2 GB for human genome) owing to aggressive lossless BWT compression, but the use of data compression increases the number of memory accesses required per data structure lookup. Bowtie2’s total alignment time is the greatest among the aligners evaluated in this work, consistent with the additional memory-access overhead introduced by its highly compressed index representation.

#### minibwa (2026)

minibwa [16] is a recent algorithm that combines BWA-MEM’s variable-length SMEM seeding strategy with minimap2’s chaining and SIMD-based base alignment. Its seeding stage combines: (a) the ropebwt3 batched SMEM-finding algorithm [26], (b) decoupling memory access from computation via software prefetch so that independent query strings can mostly hide their FM index cache miss latency behind another string’s computation, and (c) applies the same batching strategy to look up reference positions in the suffix array using the SA interval. Interestingly, while minibwa’s FM index design layout is essentially unchanged from BWA-MEM’s, its reported speedup is approximately 4× over BWA-MEM, and greater than 2× over BWA-MEM2. In our single-thread experiments using the T2T-CHM13 reference, minibwa’s peak memory was 8.3 GB and the speedup comes from hiding memory latency rather than reducing the number or cost of memory accesses per lookup. Beyond latency hiding, minibwa reduces the downstream processing cost in two ways. First, its minimap2-based chaining algorithm produces fewer unmapped mates in paired-end mode, reducing invocations of the mate-rescue procedure, a fallback step in which Smith–Waterman alignment is applied over a large reference window centered on the successfully mapped mate’s position to force-align its partner, which is one of the more expensive steps in paired-end alignment. Second, minibwa applies a hard limit of 50 chains per read, discarding all seeds beyond this threshold. This cap substantially reduces chaining and extension work for reads mapping to highly repetitive regions of the reference, where seed generation would otherwise produce a large number of candidate chains. The cost of both optimizations is reduced alignment accuracy when compared with BWA-MEM2.

#### Strobealign (2022)

Strobealign [13] precomputes a table of fuzzy, variable-length seeds. It first subsamples the reference into open *syncmers* [14]. A syncmer is a *k*-mer that is kept only when the smallest of its overlapping *s*-mers (ranked by a hash value) lies at a fixed position within the *k*- mer. This condition holds for roughly one in five *k*-mers, so only that fraction is retained. Nearby syncmer pairs are then linked into gapped *strobemer* seeds [15], in which two shorter subsequences separated by a variable gap are hashed into a single seed. Because a strobemer spans two syncmers, it attains the uniqueness of a much longer exact *k*-mer while tolerating substitutions and short indels between the two anchors, reducing seed repetitiveness on the reference. Candidate positions are retrieved by hash lookups into a sorted seed table rather than using base-by-base index traversal. The resulting matches are chained and passed to a Striped Smith–Waterman extension [27], a SIMD-vectorised formulation of Smith–Waterman in which the dynamic programming cells are laid out in a striped order so that several cells can be evaluated fast in parallel within one vector register. For read lengths of 100-250 bp, Strobealign is 4.5–6 times faster than BWA-MEM2 while providing comparable accuracy [13].

#### Summary

The alignment algorithms reviewed above collectively demonstrate that seeding speed, memory footprint, and compatibility with BWAMEM2’s downstream pipeline are in fundamental tension: (a) ERT gains speed at the cost of a larger memory footprint that is required by the relatively large ERT data structure; (b) minimizer- and strobemer-based methods reduce the memory footprint but subsample the seed space, trading off short read sensitivity (the fraction of reads for which a valid alignment is found) for speed; and

(c) recent aligners, such as minibwa, gain speed relative to BWAMEM2 but at the cost of accuracy. This challenge is particularly pronounced for the short reads generated by contemporary high-volume NGS machines, where hundreds of millions of reads must be aligned rapidly and accurately to a human reference despite the presence of large repetitious regions. RosaSeed addresses these tradeoffs through a combination of an *s*-base reference encoding and a corresponding *s*-base FM index, “faster exact” alignment of seeds using a jump table, and configurable multi-phase seeding algorithms that preserve compatibility with the downstream BWA-MEM2 seed chaining and alignment stages, as described next.

## 3. Methods Used in RosaSeed

This section describes the design, implementation and evaluation methodology of RosaSeed, focusing on its seeding stage. We first present the generalized *s*-base derived reference representation and its FM index, followed by the checkpoint-based occurrence array and jump table acceleration method used to speed up FM index traversal. We then describe the Phase A primary seeding algorithm, the supplementary seeding strategies employed by Phases B and C, and their optimization to suit the downstream BWA-MEM2 alignment pipeline. We then describe the pre-chain pruning of seeds to avoid unproductive work for the seed chaining and BSW stages. Finally, we review the lossless compression of the SA, configurable operating modes and coroutine-based prefetch scheduling, and the experimental conditions that were used for benchmarking, and the accuracy metrics that were used to compare the aligners.

### 3.1. s-base reference and FM index construction

Standard FM index implementations for read aligners operate over a 4-symbol *alphabet* Σ = {A, C, G, T}, aligning one base per backward-search extension step (alignment step) and incurring two accesses to the large occurrence array, Occ(*σ*, ℓ) and Occ(*σ, h*), per step. RosaSeed instead operates over a *radix-*4^*s*^ *alphabet* in which each symbol encodes *s* ≥ 1 consecutive DNA bases, reducing the number of FM index steps required per seed extension by a factor of *s* [28]. The recommended RosaSeed configuration proposed in this work uses *s* = 2, which encodes two adjacent DNA bases as one radix-16 symbol. The RosaSeed framework also supports *s* = 1, which corresponds to the conventional radix-4 representation, and *s* = 3, which encodes three adjacent DNA bases as one radix-64 symbol. Thus, for *s* = 1 the alphabet has 4 single-base symbols, for *s* = 2 it has 16 2-base symbols, and for *s* = 3 it has 64 3-base symbols. Compared with standard radix-4 traversal, the *s* = 2 and *s* = 3 configurations reduce the backward traversal depth to approximately one half and one third, respectively, thereby reducing the number of occurrence array accesses per seed. Our strategy was inspired by the *n*-step FM index of Chacón et al. [28] but our implementation imposes no divisibility constraints on the query length and it operates seamlessly across arbitrary read boundaries and across concatenated forward and reverse-complement reference strands.

#### Construction of the s-base reference

The same construction is used for *s* = 1, 2 and 3. Let *F* denote the forward strand of a reference of length *L*. To handle arbitrary seed starting positions, RosaSeed constructs *s* phase-shifted symbol sequences, one for each possible starting offset, which are subsequently concatenated into a single *derived reference*. For *s* = 2, each phase is encoded as a sequence of adjacent 2-base symbols. The *even phase* encodes base pairs (*F* [2*i*], *F* [2*i*+1]) for *i* ≥ 0, while the *odd phase* encodes base pairs (*F* [2*i*+1], *F* [2*i*+2]), for *i* = 0, 1, 2, …, with analogous constructions for *F* (the reverse complement strand of the reference of length *L*). Thus we have:

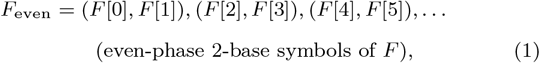

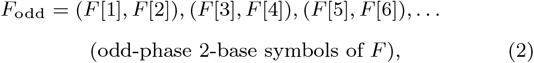

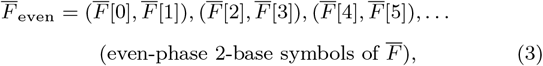

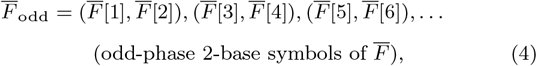

The corresponding 2-base derived reference *Ref* is then the concatenation

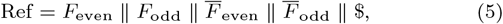

where ∥ denotes sequence concatenation and $ is the terminating symbol for the derived reference that is included in the conventional definition of the BWT. Each 2-base symbol is represented by a single radix-16 digit in the set `{0 × 0, 0 × 1 … 0xF}`. (Note that `A`–`F` denote hexadecimal digits rather than DNA bases.) Together, the phase-shifted symbol sequences *F*_even_ and *F*_odd_ (and their reverse-complement counterparts) cover every possible base-level positions in the original reference, ensuring that no seed starting position is missed, regardless of its offset in the original reference. A single FM index is then constructed in the normal way from this 2-base derived reference. Figure 3 illustrates this construction for a small example, including the 2-base derived reference *Ref*, resulting suffix array (SA), Burrows–Wheeler Transform (BWT), occurrence array (Occ), and count array (C). RosaSeed uses the open source *gsufsort* package to construct SA and BWT from the *s*-base reference during index construction [29].

**Figure 3.**
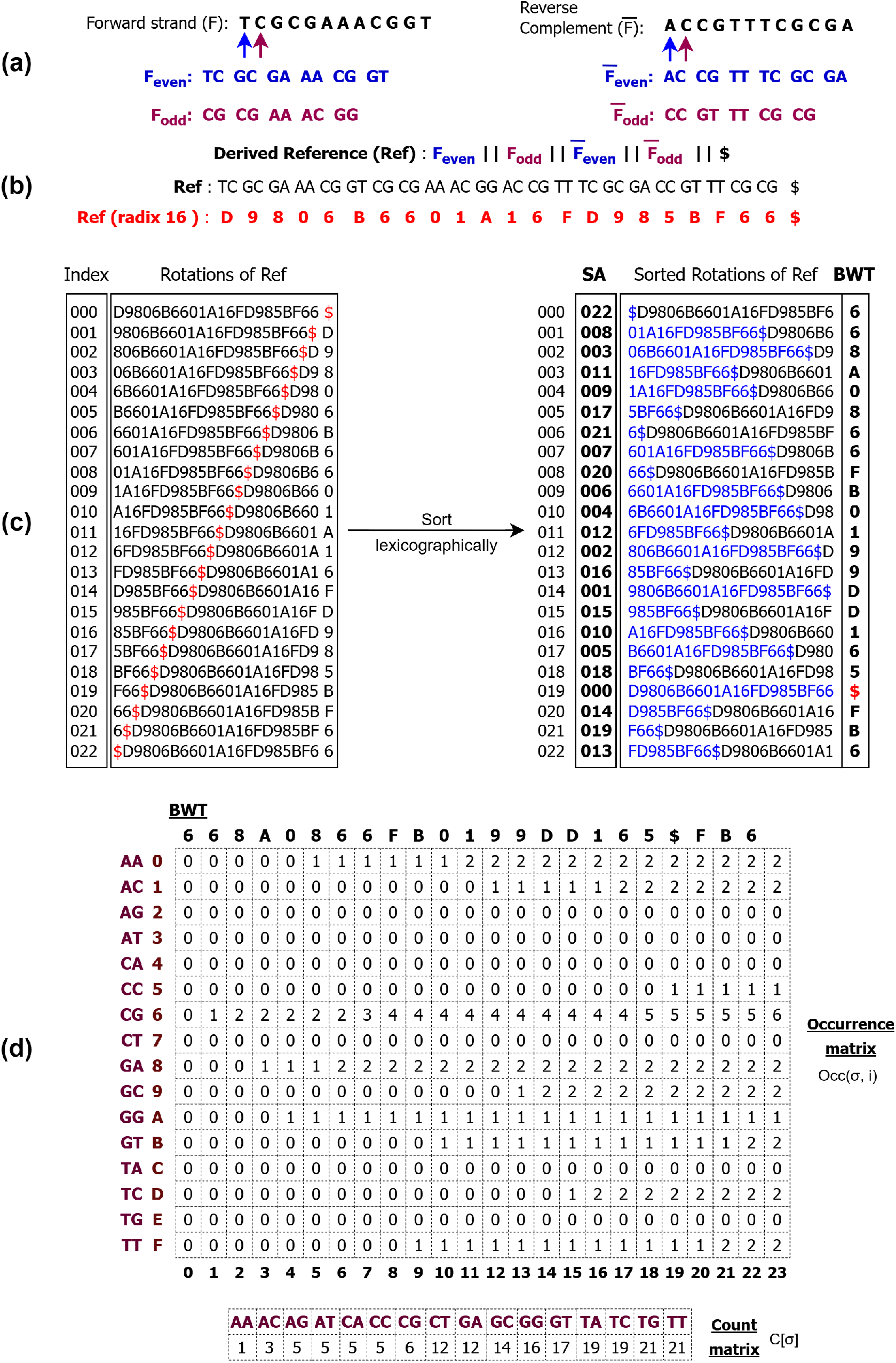
FM index construction for an *s*=2 (radix-16) derived reference. (a) Even-phase and odd-phase 2-base symbol sequences are constructed from the forward and reverse-complement strands *F* and 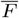. (b) The derived reference *Ref* is built in the order 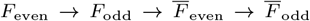 and encoded using radix-16 symbols. (c) The rows in the rotation array are lexicographically sorted to produce the suffix array (SA) and BWT. (d) The occurrence array Occ(*σ, i*) and count array *C*[*σ*] contain the information used by the FM index backward-search recurrence ℓ ← *C*[*σ*] + Occ(*σ*, ℓ), *h* ← *C*[*σ*] + Occ(*σ, h*) that updates ℓ and *h*, when the next symbol *σ* is swept.

### 3.2. The checkpoint-based compressed occurrence array

The occurrence array Occ(*σ, i*) records the number of occurrences of symbol *σ* in the BWT prefix BWT[0, *i*) = (BWT[0], …, BWT[*i* − 1]) and is consulted at every backward-search step. RosaSeed employs a checkpoint-based compressed occurrence array derived from BWAMEM2 [11] but generalized to support the multiple alphabet sizes corresponding to the RosaSeed configurations with different values of *s*. Specifically, the occurrence array supports the conventional radix-4 alphabet (*s* = 1), the radix-16 2-base alphabet (*s* = 2), and the radix-64 3-base alphabet (*s* = 3). Similar to the compressed SA that can be used in BWA-MEM2, checkpoints are formed and stored at fixed intervals along the BWT [11]. Each checkpoint stores cumulative symbol counts up to the checkpoint boundary together with a compact encoded representation of the subsequent BWT block. In Figure 3, the row labels in panel (d) carry a dual notation: the left part identifies the 2-base pair (e.g. `CG`) and the right part gives its radix-16 hexadecimal encoding (e.g. `6`). The 24 column indices *i* = 0, 1, …, 23 correspond to query positions in the exclusive prefix count Occ(*σ, i*) = |{*j < i* : BWT[*j*] = *σ*}|, so column *i* = 0 is identically zero for every symbol, and column *i* = 23 equals the total count of *σ* in the entire BWT. Each entry increases by exactly one relative to its left neighbour when BWT[*i* − 1] = *σ*, and remains unchanged otherwise. For example, the row for `CG` (*σ* = 6) increments at columns *i* = 1, 2, 7, 8, 18, 23 because BWT[0] = BWT[1] = BWT[6] = BWT[7] = BWT[17] = BWT[22] = 6, yielding the final count of six. During backward search, an occurrence query is answered by locating the nearest preceding checkpoint and then computing the contribution of symbols between the checkpoint and the query position using bit-parallel population count operations. Because backward search requires two occurrence array lookups per extension step, reducing the number of extension steps through larger radix alphabets (*s* = 2) directly reduces the total number of occurrence array accesses performed during seeding. Also, since each checkpoint stores cumulative counts for every symbol in the alphabet Σ, the memory footprint of the occurrence array depends on both the alphabet size |Σ| and the checkpoint sampling interval. Specifically, the storage required per checkpoint scales with the alphabet size |Σ| = 4^*s*^, while the total number of checkpoints is inversely proportional to the sampling distance. Consequently, using larger radix alphabets (*s* = 2 and *s* = 3) and more frequent checkpointing with smaller intervals increases the size of the occurrence array, whereas using larger sampling intervals reduces the memory footprint at the cost of additional symbol processing during occurrence queries.

### 3.3. The jump table

A *jump table (JT)* is a data structure that is precomputed from the derived reference and that stores the suffix array interval corresponding to every possible *k*-mer prefix of some fixed length *k >* 1 over the radix-4^*s*^ alphabet. Rather than performing the initial sequence of *k* FM index backward search extensions, RosaSeed begins seed generation with a single jump table lookup, for the rightmost *k*-mer in the next portion of the read, that returns the corresponding suffix array interval, thereby significantly reducing the number of occurrence array accesses that would otherwise be required.

The idea of precomputing suffix array intervals for short prefixes (*k*-mers) has been utilized previously: for example by Ning et al. [30] for rapid database sequence querying, and by Arram et al. [31] for FPGA-accelerated sequence alignment. RosaSeed adapts this concept to a software CPU implementation where the jump table entries are encoded differently (explained below) and extends it to support multiple values of *s* corresponding to the one-, two-, and 3-base symbol alphabets, enabling JT acceleration across different radix-4^*s*^ FM index configurations.

For clarity, the *jump table length* refers to the number of bases in the first JT index and not the number of radix-4^*s*^ symbols. The JT therefore contains 4^*k*^ entries, one for each possible base-level *k*-mer. Internally, for an *s*-base index value, this *k*-mer is interpreted as a sequence of radix-4^*s*^ symbols where applicable. Each entry in the JT stores the SA interval [ℓ, *h*) associated with the corresponding prefix, where ℓ and *h* denote the *lower* and *upper SA boundaries*, respectively. The *interval size* is defined as *d* = *h* − ℓ, corresponding to the number of suffixes contained in the interval, which we will call the *abundance* of the associated *k*- mer in the reference genome. Consequently, the jump table replaces the initial *k*-base prefix traversal with a single memory lookup. For the primary 2-base RosaSeed design (*s* = 2), the effects of JT lengths corresponding to *k* ∈ {14, 15, 16} bases were evaluated experimentally. These configurations require approximately 2 GB, 8 GB, and 32 GB of memory, respectively, when using the 8-byte entry representation employed in RosaSeed. Larger jump tables reduce the amount of FM index traversal required during seeding at the expense of an increased JT memory footprint.

RosaSeed employs a 64-bit JT encoding that avoids SA accesses for the relatively common case of a unique SA interval. Bit 63 is a uniqueness flag. When the flag is 0, the entry represents a non-unique interval and it stores the lower SA boundary ℓ in the lower 33 bits together with the interval width *d* = *h* − ℓ in the remaining 30 bits. When the flag is 1, the entry corresponds to a *unique interval* (*d* = 1) and it directly stores the reference position in the lower 63 bits. Thus, prefixes whose SA interval width is one can be resolved quickly with a single JT lookup without requiring any suffix array accesses. Consequently, suffix array decompression is required only for nonunique JT entries, where reference positions must be recovered from sampled suffix array values using a Last-to-First (LF) [12] mapping.

### 3.4. Phase A: strand-aware disjoint pivot sweep

Phase A is the proposed new initial seeding stage shared by all RosaSeed configurations. Starting at the rightmost read position, *p* = *N* −1, pivot positions in the read are considered sweeping from right to left. At each pivot, a seed candidate is produced from a JT lookup followed by zero or more *s*-base backward extension steps. Candidate seeds that satisfy the Phase A emission criteria are added (that is, *emitted*) to the *output seed list S*_*A*_. List *S*_*A*_ is passed to either the Phase B or Phase C stage, depending on the selected RosaSeed configuration. The high-level Phase A control flow is shared across the *s*-base RosaSeed designs, as shown in Figure 4.

**Figure 4.**
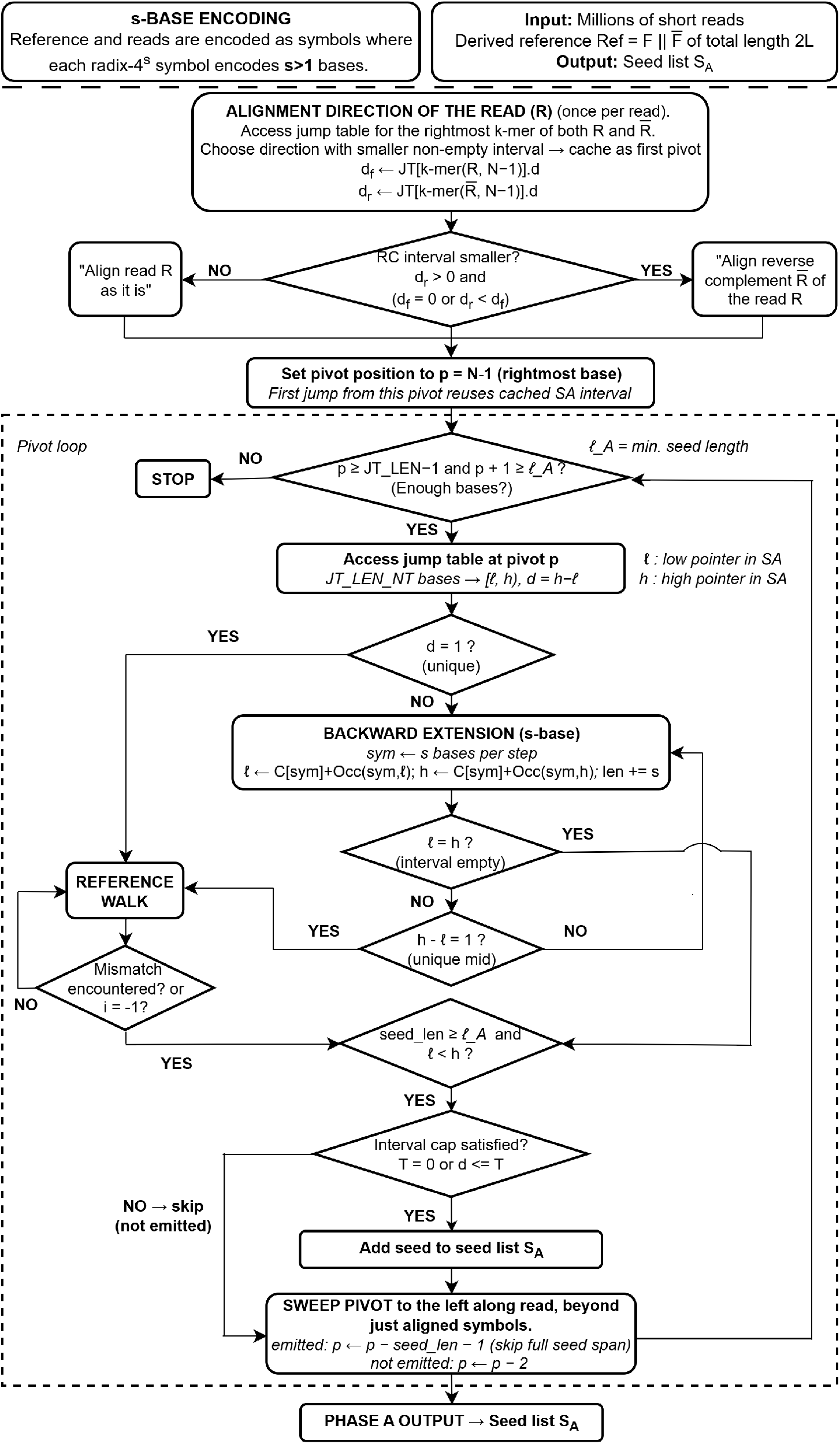
Phase A control flow. *Top*: The alignment direction selection heuristic accesses the jump table for the rightmost *k*-mer on the read and its reverse complement (RC), thus determining SA interval widths *d*_*f*_ (forward) and *d*_*r*_ (RC). The RC of the read is selected for processing if *d*_*r*_ *>* 0 and (*d*_*f*_ = 0 or *d*_*r*_ *< d*_*f*_); otherwise the read is used as is. Phase A performs a right–to–left pivot sweep over the chosen read strand. *Dashed box (Pivot loop)*: at each pivot position *p*, a jump table lookup retrieves the SA interval [ℓ, *h*) with *d* = *h*−ℓ. Unique hits (*d* = 1) immediately trigger a reference look up; non-unique hits cause an *s*-base backward extension (ℓ ← *C*[*σ*] + Occ(*σ*, ℓ), *h* ← *C*[*σ*] + Occ(*σ, h*)) until either the interval empties, becomes unique, or the opposite end of the read is reached. Seeds passing the minimum length filter (seed len ≥ ℓ_*A*_ = 19 bases) and maximum SA interval cap (*T* = 0 or *d* ≤ *T*) heuristics are emitted and the next pivot is placed at *p* − seed len − 1 (disjoint); otherwise, one base is skipped, and the next pivot is placed two positions to the left of the current pivot. *Bottom*: *S*_*A*_ is passed to Phase B or Phase C, depending on the RosaSeed configuration.

#### Alignment Direction

Before the pivot sweep along a read *R* is started, RosaSeed accesses the JT once at position *p* = *N* −1 for both the *R* and its reverse complement 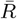, yielding two SA interval widths:

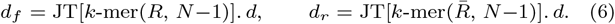

The RC of the read is selected as the preferred direction if *d*_*r*_ *>* 0 and either *d*_*f*_ = 0 or *d*_*r*_ *< d*_*f*_ ; otherwise, the forward read (read as is) is used. We call this heuristically selected representation of the read, *R* or 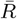, as *pat*. The selected intervals [ℓ, *h*) are cached to avoid a redundant jump table lookup at the first pivot. This two-probe heuristic costs one additional JT access per read, but by selecting the read strand associated with the smaller initial SA interval, the size of the expected search space for subsequent extension has been found experimentally to be reduced on average.

#### Pivot loop

The pivot processing loop iterates while *p* ≥ *k* − 1 and *p* + 1 ≥ ℓ_*A*_, where *k* is the *k*-mer length for the jump table and ℓ_*A*_ = 19 bases is the minimum acceptable seed length, using the same criteria as BWA-MEM2. At each pivot, three sequential stages are executed.

#### Jump table lookup

The precomputed SA interval of the k-mer [ℓ, *h*) with *d* = *h* − ℓ is retrieved with one JT look up. For the first pivot of the *pat*, the SA interval that was cached during the alignment direction selection, is reused.

#### Seed extension

One of two paths is taken based on the SA interval distance *d*:

- **Unique at jump** (*d* = 1): The reference position is extracted from the lower 63 bits of the 8-byte JT entry. A *reference walk* then extends the seed leftward along the reference genome, matching symbols against *pat* one encoded symbol at a time (for the recommended baseline configuration, RosaSeed, two bases per symbol) until either a mismatch is encountered or the left edge of the read is reached.
- **Non-unique** (*d >* 1): *s*-base backward extension proceeds via the recurrence

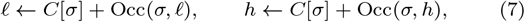

where *σ* is the radix-4^*s*^ symbol that encodes the next *s* read bases ending at position *i*. For a 2-base RosaSeed design, this symbol is the radix-16 encoding of (*pat*[*i*−1], *pat*[*i*]). Three stopping conditions terminate the loop: (i) the SA interval empties (ℓ = *h*) and the step attempt is rolled back; (ii) the SA interval becomes unique (*h* − ℓ = 1) and a reference walk is triggered; (iii) the left edge of the read is reached. Phase A does *not* apply an early-stop based on the interval size: it always seeks the *longest possible seed* from each pivot.

#### Seed emission

A seed is emitted to the output list if and only if all three of the following conditions hold:

1. **The SA Interval is non-empty**: *d* = *h* − ℓ *>* 0
2. **Length filter (as in BWA-MEM2)**: seed len ≥ ℓ_*A*_(19 nt)
3. **Phase A SA-interval cap T**: either *T* = 0 (disabled) or the SA interval *d* ≤ *T*

Seeds with *d > T* correspond to *k*-mers present at more than *T* locations in the reference.

For example, *T* = 5,000, BWA-MEM2’s downstream chaining stage processes at most 500 alignment locations per seed, by default. When *d* is much larger than this SA interval cap, only a small fraction of candidate locations can be examined, reducing the likelihood that such repetitive (that is, abundant) seeds contribute useful alignment information. Abundant seeds with large SA intervals can therefore be suppressed relatively safely during Phase A, with the resulting coverage gaps filled in by Phase C (Section 3.6). Setting *T* = 0 disables the SA interval cap, thus allowing all seeds meeting the minimum length filter to be emitted to the seed chaining phase regardless of the SA interval size *d*.

#### Pivot advance and output

If a seed meets the three emission criteria, the next pivot is placed immediately to the left of the emitted seed after skipping one intervening base. Equivalently, for a seed ending at pivot *p* with length *seed len*, the next pivot is positioned at *p* − *seed len* − 1. This ensures that successive Phase A seeds are disjoint and that the base immediately preceding the emitted seed is not reconsidered. If no seed is emitted, Phase A skips the base immediately preceding the current pivot and resumes at *p* − 2, allowing the next JT look-up the opportunity to identify an entirely new starting *k*-mer.

Phase A produces a seed list *S*_*A*_, sorted by start position in the forward read strand position index system, before the supplementary stage (either Phase B or C) is invoked. For reads mapped onto the RC read strand, the seed positions are mapped back to the forward read strand index system before insertion into *S*_*A*_.

### 3.5. Phase B: fixed-pivot supplementary seeding

When enabled at compile time, Phase B provides supplementary seeding beyond what was accomplished in Phase A by placing three fixed pivots (user-configurable via the `-pb` setting; the default is 3 for *N* =150 bp reads) at evenly spaced positions along the read:

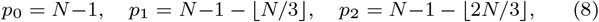

where *p* ∈ {149, 99, 49} for *N* = 150 nt reads. Phase B adopts the supplementary seed emission criterion used by BWA-MEM2’s low-interval seeding strategy but evaluates it only at a small number of fixed pivots using RosaSeed’s *s*-base FM index and jump table. The high-level Phase B control flow is shared across the *s*-base RosaSeed variants, as illustrated in Figure 5 for the default case of `-pb`=3.

**Figure 5.**
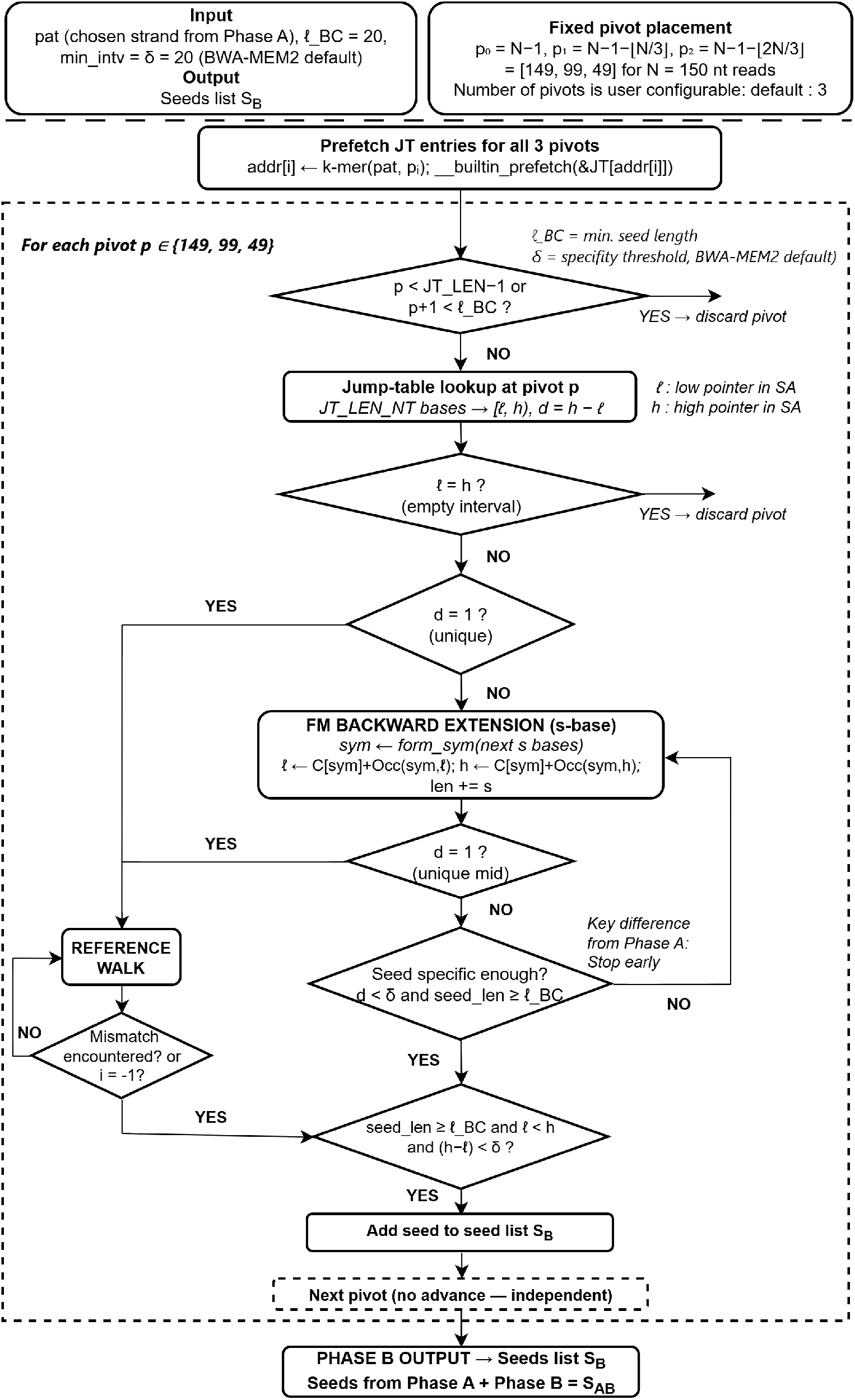
Phase B control flow. *Top*: Phase B receives the chosen read strand pattern *pat* from Phase A and identifies the three pivot positions in the read *{*149, 99, 49*}* for *N* = 150 nt reads. Jump table entries for all three pivot *k*-mers are prefetched before the pivot loop to avoid repeated jump table accesses during per-pivot processing. *Pivot loop (dashed box)*: Each pivot is processed independently. Heuristic checks cause a pivot to be skipped if *p < k* − 1 or *p* + 1 *<* ℓ_*BC*_. Unique hits (*d* = 1) trigger a reference walk; non-unique hits undergo *s*-base FM backward extension (Equation (7)). If an interval becomes unique during backward extension, Phase B transitions to a reference walk. *Early stop (coral diamond)*: Unlike Phase A, Phase B stops seed extension as soon as *d < δ* and seed len ≥ ℓ_*BC*_ – the key algorithmic difference. The emission condition requires seed len ≥ ℓ_*BC*_, ℓ *< h*, and (*h*−ℓ) *< δ. Bottom*: The Phase B output is merged with *S*_*A*_ to form *S*_*AB*_, and then passed to the BWA-MEM2 chaining phase. The parameter settings: ℓ_*BC*_ = 20 nt and *δ* = 20 match the default supplementary seed length and interval width thresholds used in BWA-MEM2.

#### Per-pivot processing

In phase B each pivot *p*_*i*_ is processed independently. A pivot is skipped if either *p*_*i*_ *< k* − 1 (insufficient bases for a JT query) or if *p*_*i*_ + 1 *<* ℓ_*BC*_ (insufficient bases were aligned to meet the minimum seed length ℓ_*BC*_ = ℓ_*A*_ + 1 = 20 bases for Phase B).

Extension follows the same two paths as in Phase A: (1) a reference walk for unique hits (to rapidly confirm a likely unique alignment location in the reference), or (2) an *s*-base backward extension, following Equation 7 for non-unique hits. If a non-unique interval becomes unique (that is, *h* − ℓ = 1) during backward extension, Phase B transitions to a reference walk, as in Phase A. However, Phase B also applies the following *early-stop* condition:

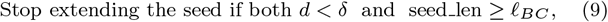

where *δ* is the SA interval width threshold (corresponding to BWAMEM2’s `min_intv` parameter). A seed is emitted if the three conditions seed len ≥ ℓ_*BC*_, ℓ *< h*, and (*h* − ℓ) *< δ* are met. Since Phase B operates on only a relatively small fixed number of pivots, its objective is to rapidly identify additional specific seeds at strategically distributed locations across the read. Once the interval drops below *δ* and the minimum seed length ℓ_*BC*_ is met, seed extension stops in Phase B. After processing each pivot (whether or not a seed is emitted), the algorithm moves to the next pivot (if present) in Phase B. Note that a Phase B seed may be identical to a seed already present in *S*_*A*_.

#### Output and comparison with Phase A

Phase B produces up to three additional seeds (`-pb`: the default is 3). The combined seed list *S*_*AB*_ consists of seeds emitted by Phase A and Phase B. Phase B uses BWA-MEM2’s supplementary seed thresholds for the minimum seed length and SA interval width, denoted here as ℓ_*BC*_ and *δ*, respectively. Phase B complements the disjoint Phase A sweep by improving the seed coverage through a small number of spaced-out supplementary pivots while requiring only a limited number of additional *s*-base traversals. Table 1 summarizes the key algorithmic differences between Phases A and B.

**Table 1.** Key algorithmic differences between Phase A and Phase B.

| Property | Phase A | Phase B |
| --- | --- | --- |
| Considered pivot positions | Adaptive right→left pivot sweep | User configurable; default = 3:<br>$\{N - 1, N - 1 - \lfloor N/3 \rfloor, N - 1 - \lfloor 2N/3 \rfloor\}$. |
| Extension goal | Longest possible seed | Stop once a sufficiently specific seed ($d < \delta$) is obtained. |
| Early-stop condition | None; extends to failure | Stop when $d < \delta$ and $\text{seed\_len} \geq \ell_{BC}$. |
| SA-interval criterion | Optional SA-interval cap: $d \leq T$ | Maximum interval-width threshold: $(d = h - \ell) < \delta$. |
| Minimum seed length | $\ell_A$ nt | $\ell_{BC}$ nt |
| Seed overlap | Disjoint by construction within $S_A$ | May overlap with $S_A$ or other Phase B seeds. |

Unlike Phase A, which aims to find the longest possible disjoint seeds through exhaustive extension from every pivot, Phase B prioritizes computational efficiency by terminating seed extension as soon as a sufficiently specific supplementary seed has been identified at a small number of strategically selected pivots.

### 3.6. Phase C: Adaptive Gap-Directed Supplementary Seeding

#### Motivation

Unaligned intervals, called *gaps*, can arise anywhere along the read: at the left end, where the right-to-left Phase A sweep must terminate; in the interior, where the Phase A skips over a region that no pivot reaches; or at the right end, for cases where the first pivot fails to produce a qualifying seed. Although BWA-MEM2’s BSW extension stage can postulate mutations that allow adjacent seeds in a chain to be joined, it cannot compensate for the absence of seeds that would fill a longer gap. BSW operates only on candidate regions of exact alignment specified by seeds that are found during seeding and grouped during chaining: A nonaligned read region unsupported by any seed will not produce a seed chain of sufficiently high score to survive filtering and thus trigger BSW extension during local alignment. The fixed pivots used in Phase B provide a mechanism for finding seeds that increase the alignment coverage in the read; however, uncovered intervals vary in size and position across reads, motivating a supplementary adaptive seeding strategy that places additional pivots directly within encountered alignment gaps.

Phase C is a gap-directed supplementary seeding strategy that inspects the sorted Phase A seed set *S*_*A*_, identifies unaligned gaps along the read, and evaluates additional pivots within each gap using the jump table and *s*-base FM index already resident in memory. Since these supplementary seeds are generated by examining unaligned gaps between Phase A seeds, we call them *gap-directed* seeds. Unlike Phase B, whose pivot locations are fixed a priori, Phase C adapts its pivot placement to the observed seed distribution from Phase A for each read. While Phase C does not guarantee that considering every gap will produce a qualifying seed, it systematically examines the gaps and increases the likelihood of finding additional seeds that satisfy the minimum seed length and specificity thresholds. This gap-directed strategy is independent of the underlying symbol encoding and is therefore applicable to all of the 1-, 2- and 3-base versions of RosaSeed.

Figure 6 illustrates the motivation for gap-directed supplementary seeding. Phase A generates non-overlapping seeds that leave unaligned intervals (yellow) between successive seeds. Although the fixed supplementary pivots (used in Phase B) can improve the seed coverage, the locations and sizes of the gaps left by Phase A are sequence-dependent and cannot be predicted a priori.

**Figure 6.**
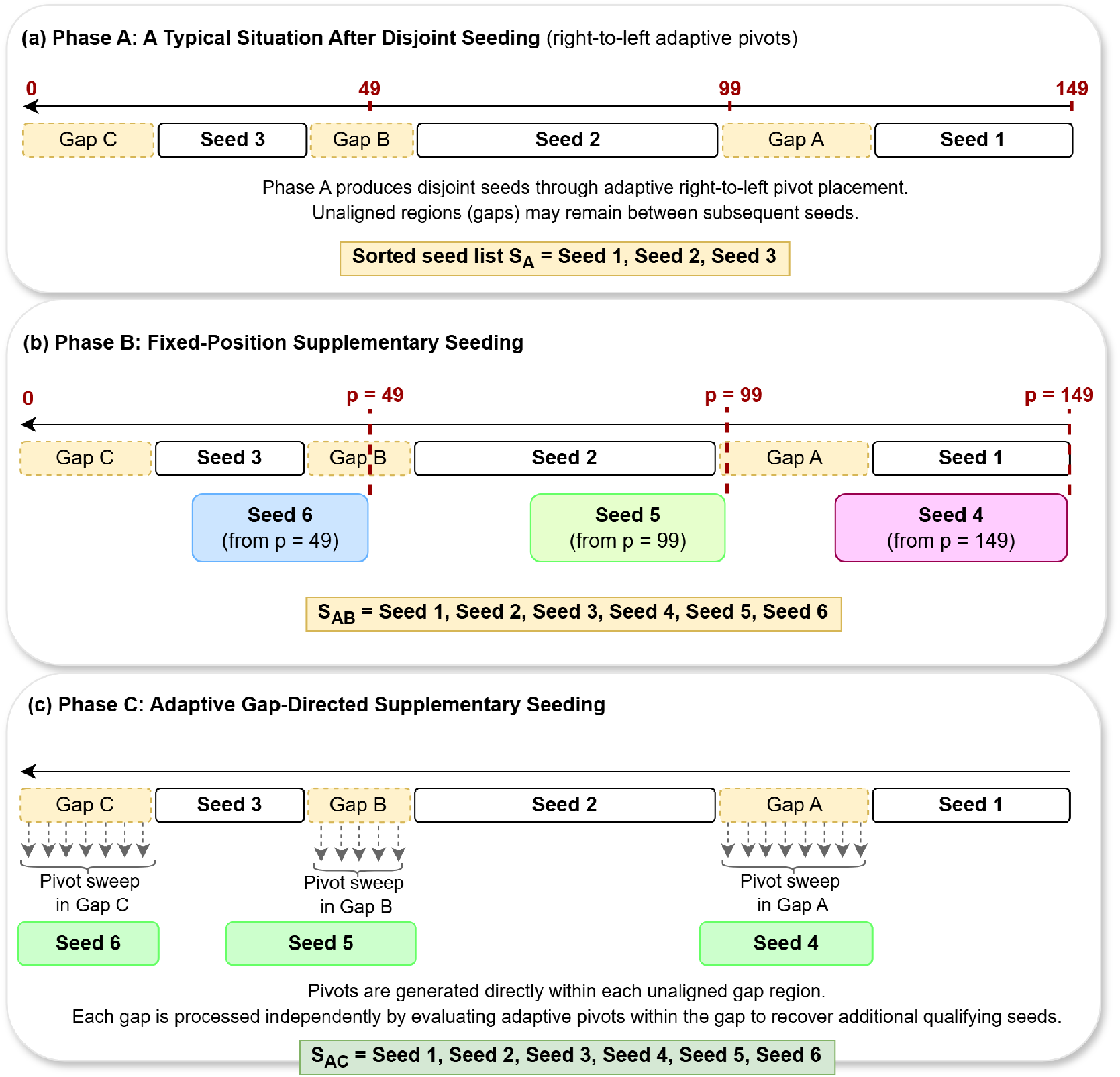
Motivation for adaptive gap-directed supplementary seeding. (a) Phase A performs an adaptive right-to-left disjoint pivot sweep, producing a primary seed set *S*_*A*_ that may leave unaligned intervals (gaps) between adjacent seeds. (b) Phase B supplements *S*_*A*_ by evaluating a small number of fixed, evenly spaced pivots, that are evaluated to generate more seeds. (c) Phase C supplements *S*_*A*_ by adaptively evaluating pivots within gaps identified after Phase A. Phase B and Phase C are complementary supplementary seeding strategies that differ in their pivot-selection mechanism while sharing the objective of improving seed coverage beyond that of Phase A.

#### Gap Identification

Phase C inputs the Phase A seed list *S*_*A*_ and then sorts it according to the forward-read coordinate system before gap identification is performed. Phase C then identifies three categories of unaligned gaps in the read:

- *Left-edge*: [0, *S*_*A*_[0].start − 1].
- *Internal* : [*S*_*A*_[*i*].end + 1, *S*_*A*_[*i*+1].start − 1] for each consecutive pair of seeds.
- *Right-edge*: [*S*_*A*_[last].end + 1, *N* − 1]. *S*_*A*_[0].start−1 points to the position in the *pat* that is exactly one base (or index) before the beginning of the first seed. If *S*_*A*_ is empty, the entire read interval [0, *N* − 1] is treated as a single gap. Gaps smaller than a setting Γ nt (such as Γ=5) are skipped, as their omission has been found experimentally to have a negligible effect on chaining.

#### Adaptive Pivot Placement

For a gap *G* = [*g*_start_, *g*_end_] of size *g* = *g*_end_ − *g*_start_ + 1, Phase C constructs a pivot list by stepping right-to-left across the gap:

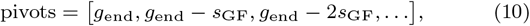

where *s*_GF_ denotes the gap-directed pivot spacing. For short gaps (*g* ≤ ℓ_*BC*_, for a settable value ℓ_*BC*_), a denser spacing of max(1, ⌊*s*_GF_*/*2⌋) is used, whereas larger gaps use the nominal spacing *s*_GF_ (also user-configurable). This allows Phase C to explore small gaps more aggressively while avoiding excessive pivot evaluations in larger gaps. Each pivot is processed using the same JT lookup, reference walk, and *s*-base FM index extension procedures used in Phases A and B. Each generated pivot is evaluated at most once. Phase C then advances to the next pivot in the gap until all candidate pivots have been examined or an early-stop condition becomes true, whereupon gap processing is terminated.

#### Gap-Directed Seed Generation

For each pivot generated within a gap, Phase C first performs a JT lookup to obtain the corresponding SA interval. If the seed is unique (SA interval size *d* = 1), then seed extension is attempted with a reference walk. Otherwise, the seed is extended using the same *s*-base FM index backward extension procedure employed by Phases A and B. Extension continues until either the SA interval becomes unique, the SA interval goes empty, or the end of the read is reached.

Like Phase B, Phase C is specificity-driven (by seeking seeds with smaller abundance) rather than seed-length driven. Seed extension stops once the SA interval size falls below a settable threshold *δ* (by default *δ* = 20) and the minimum acceptable seed length ℓ_*BC*_ has been reached.

#### Gap Coverage and Strand-Specific Recovery

Phase C first evaluates gap-directed pivots on the pat selected during Phase A, which may be either the forward read or its reverse complement. When the compile-time option `GAPFILL_ALWAYS_RUN_BOTH_STRANDS` is disabled, seeding is performed only along the heuristically selected alignment direction, *pat*. When the option is enabled, as in the recommended RosaSeed configuration, the corresponding pivot set is evaluated on both read strands regardless of the outcome of the first pass. Any seeds emitted from the reverse complement read are mapped back to the forward-direction reference coordinates before being added to the output seed list.

After seed emission, Phase C evaluates whether the current gap has been covered. A gap *G* = [*g*_start_, *g*_end_] is considered covered (aligned) only when at least one emitted seed spans the entire gap interval; partial overlap with the gap is not sufficient. Formally, coverage requires

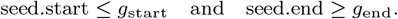

When the compile-time option `GAPFILL_EARLY_EXIT` is enabled, pivot evaluation terminates as soon as the current gap is covered, thereby reducing the number of JT lookups and FM index extensions performed for that gap. This greedy early-exit strategy was found to reduce computational overhead while preserving alignment accuracy in the evaluated configuration.

It is to be noted that in the recommended RosaSeed, both `GAPFILL ALWAYS RUN BOTH STRANDS` and `GAPFILL EARLY EXIT` are enabled. Early exit therefore applies independently within each read strand-specific pass: covering the gap during the pass on pat terminates further pivot evaluation for that pass, but does not suppress the subsequent pass on the opposite read strand.

#### Interaction with the Phase A SA-interval Cap

Phase A optionally applies a maximum SA interval cap *T* to prune away seeds whose abundance exceeds a user-specified threshold. Such seeds are typically mapped to highly repetitive regions of the reference and thus align to a large number of reference locations, making them less informative (and more time-consuming) for downstream chaining and BSW-based alignment.

Suppressing a seed because the SA interval size *d > T* may leave an uncovered gap in the Phase A seed set *S*_*A*_. These uncovered gaps subsequently become candidate gaps for Phase C. Rather than forwarding a highly abundant seed to the chaining stage, RosaSeed attempts to find additional seeds from the same read region by evaluating gap-directed pivots within the resulting gap.

The success of this supplementary seeding strategy depends on the local sequence context. In matches to moderately repetitive regions of the reference, Phase C often finds shorter but more specific seeds that satisfy the supplementary seeding criterion (*d < δ*) and therefore contribute to useful chaining. However, in matches to highly repetitive regions of the reference, no qualifying seed may exist regardless of pivot placement. Consequently, the interval cap and Phase C operate as complementary mechanisms: The SA interval cap reduces the propagation of very abundant seeds that are unlikely to contribute useful chaining information, while Phase C attempts to find more specific, and hence informative, seeds from the resulting gap regions.

#### Comparing the Phase B and Phase C seeding strategies

Phases B and C represent two distinct supplementary seeding strategies (see Table 2). Phase B uses a small number of rapidly selected fixed pivots with minimal overhead whereas Phase C adaptively places pivots according to the alignment gaps left by Phase A, and therefore concentrates computation only on unaligned regions. Section 4 evaluates both approaches experimentally and discusses the resulting trade-offs in memory, alignment accuracy and runtime performance. Unless otherwise stated, the recommended RosaSeed configuration corresponds to the Phase A + Phase C seeding workflow. Phase B and C are mutually exclusive because both target the same goal of generating additional seeds: applying them together would generate largely overlapping seeds at the combined cost of both passes, with little additional coverage.

**Table 2.** Key algorithmic differences between the supplementary seeding strategies of Phase B and Phase C.

| Property | Phase B | Phase C |
| --- | --- | --- |
| Supplementary strategy | Fixed-position seeding | Gap-directed seeding |
| Pivot placement | Predetermined positions | Generated within uncovered gaps |
| Pivot locations | Three fixed pivots (default: right, middle, and left-third positions) | Determined dynamically from gaps left after Phase A |
| Coverage objective | Add supplementary pivots | Adaptively recover seeds from uncovered regions |
| Early-stop criterion | $d < \delta$ and $\text{seed.len} \geq \ell_{BC}$ | Same criterion |
| Read strand evaluation | Selected strand only | Selected strand with optional opposite-strand recovery |
| Interaction with Phase A interval cap | None | Attempts recovery from gaps introduced by the Phase A SA-interval cap |

### 3.7 Integration with BWA-MEM2 downstream processing

The preceding sections described the data structures and seeding algorithms used by RosaSeed to generate promising, exactly aligned seeds. These seeds must then be expressed in the representation expected by BWA-MEM2 and passed along to its downstream alignment pipeline. RosaSeed therefore looks up reference alignment positions from the SA intervals produced during seeding, constructs BWA-MEM2-compatible seed records, and forwards the resulting seed sets to the BWA-MEM2’s chaining stage. RosaSeed also applies pre-chain filtering heuristics to prune away overly abundant seeds from the generated seed set before chaining to reduce the propagation of low-value seeds. Beyond seed generation and these optional filtering steps, RosaSeed retains the original BWA-MEM2 downstream workflow, including chain construction, chain filtering, chain selection and BSW alignment extension.

#### Seed generation and reference position recovery

RosaSeed generates seeds using the derived *s*-base reference position system described in Section 3.1. Before these seeds are passed to the downstream BWA-MEM2 pipeline, RosaSeed converts the corresponding reference positions back to the corresponding baselevel reference positions. After reference position recovery, RosaSeed constructs seed records using the same logical fields expected by BWA-MEM2, including the read identifier, query start position, reference start position, seed length, and suffix array interval size. The resulting seed set is then forwarded to the standard BWA-MEM2 chaining pipeline.

#### Pre-chain singleton suppression filter

RosaSeed optionally applies a pre-chaining seed filtering heuristic designed to reduce the creation of low-value singleton (single-seed) chains before the BWA-MEM2 chaining stage. The motivation for this heuristic is to avoid letting a small number of reads generate a large number of single-seed chains originating from short and often highly abundant seeds. RosaSeed applies this heuristic at the chain construction boundary, before a candidate seed is allowed to initiate a new singleton chain. Rather than allowing every seed candidate to initiate a new singleton chain, RosaSeed may suppress candidate seeds from initiating new singleton chains when they satisfy predefined weakness criteria that can incorporate properties such as seed length and, optionally, seed abundance.

Two variants of pre-chain seed filtering are supported. The first filter suppresses the creation of new single-seed chains when the number of chains already generated for a read exceeds a predefined threshold and the candidate seed length falls below a specified minimum. The second filter additionally incorporates seed abundance information and suppresses only those short candidate seeds whose SA interval size exceeds a minimum abundance threshold before they can initiate new singleton chains. Consequently, abundance-aware seed suppression preferentially prunes highly abundant seeds while retaining short seeds that occur at relatively few reference locations.

This pruning heuristic is optional and operates identically across all RosaSeed configurations. The threshold values used in this study were selected empirically based on analyses of chain distributions and seed-abundance statistics, as described in Section 4 and Appendix table A3. Consequently, RosaSeed functions as a drop-in replacement for the BWA-MEM2 seeding stage while preserving the original downstream chaining, chain filtering, chain selection and BSW extension pipeline. The only modifications beyond seed generation are the pre-chain filtering heuristics introduced in this work to reduce the propagation of low-value seeds into downstream processing.

### 3.8. Suffix array compression

We now describe the index compression scheme that governs the memory–performance trade-off across all RosaSeed configurations. Lossless sampled SA compression with Last-to-First mapping (LF) decompression is a standard technique that was introduced in the original FM index [12] and also employed by BWAMEM2 [11]. RosaSeed supports SA compression factors of CF = 2^*x*^, where *x* ∈ {0, 1, 2, 3}, selected as a compile-time parameter (`-DSA COMPRESSION FACTOR POWER=`*x*). Only every CF-th suffix array row is stored (i.e., the sampled rows); unsampled rows are recovered by walking backward through the BWT via LF-mapping until the preceding sampled row is reached. SA compression does not affect seeding accuracy since it only changes how reference positions are resolved for non-unique seeds after their SA intervals are found. Unique JT entries store reference positions directly and therefore entirely bypass suffix array access. Consequently, SA decompression is required only when resolving reference positions associated with seeds in non-unique SA intervals.

For the recommended RosaSeed configuration, the SA when compressed by a factor of 2, occupies 14.52 GB of memory. Because RosaSeed speeds FM index traversal through 2- and 3-base symbols as well as JT acceleration, SA decompression constitutes a larger fraction of the remaining seeding cost as the compression factor increases, making optimizing the memory-performance trade-off particularly important.

### 3.9. RosaSeed operating modes & configurable parameters

RosaSeed is implemented as a configurable seeding framework that supports different *s*-base symbol alphabets, supplementary seeding strategies, SA compression, and pre-chain seed filtering heuristics. The 1-, 2- and 3-base RosaSeed configurations denote radix-4, radix-16, and radix-64 symbol encodings, respectively. These modes share the same high-level seeding pipeline but differ only in the number of DNA bases encoded per FM index symbol and, therefore, in the number of FM index extension steps required per seed.

Table 3 summarizes the principal RosaSeed configurations and algorithmic parameters that were evaluated. Parameters, such as the minimum seed length and gap-directed spacing, are reconfigurable, however, in the experiments reported below we used the default values described in the corresponding algorithm sections unless otherwise stated. SSF denotes the weak *singleton suppression filter* described in Section 3.7. SSF+A denotes the abundance-aware variant of the SSF that additionally incorporates seed abundance (SA interval size) during suppression decisions.

**Table 3.** Principal RosaSeed operating modes and configurable parameters evaluated in this work.

| Category | Options evaluated | Purpose |
| --- | --- | --- |
| Symbol multiplicity | 1-, 2-, and 3-base reference encodings | Encodes 1, 2 or 3 DNA bases per FM index symbol, corresponding to radix-4, radix-16 and radix-64 alphabets. |
| Jump table length, $k$ | $k = 14, 15$, or 16 bp where supported | Controls the length $k$ of the initial exact lookup used before FM index extension. Larger tables reduce the traversal time but increase storage cost. |
| Supplementary seeding strategy | Phase B or Phase C (default); never used together | Phase B uses fixed pivots, whereas Phase C adaptively places pivots within uncovered gaps left after Phase A. |
| Phase B pivot count | User-configurable; default is 3 | Controls the number of fixed supplementary pivots evaluated by Phase B. |
| Phase C gap-directed policy | Adaptive opposite read strand evaluation; forced opposite read strand evaluation (used in RosaSeed); optional early exit after full gap coverage (used in RosaSeed) | Controls how aggressively Phase C evaluates gap-directed pivots and whether the opposite read strand is evaluated only when needed or always. |
| Phase A SA interval cap | Default of 2000; a user-specified threshold | Suppresses highly repetitive Phase A seeds whose suffix array interval exceeds the selected cap. |
| Suffix array compression | $CF = 2^x$, $x \in \{0, 1, 2, 3\}$ | Reduces the suffix array storage cost by storing sampled SA entries and recovering unsampled entries by LF-mapping. |
| Pre-chain singleton seed suppression | SSF by default; SSF+A; can also be disabled | Suppresses weak singleton-chain seed candidates before chain construction, optionally restricting suppression to highly abundant candidates. |
| Minimum seed length | $\ell_A$ for Phase A (default 19); $\ell_{BC}$ for Phases B and C (default 20) | Sets the minimum seed length required for emission from the primary and supplementary seeding stages. |
| Gap-directed spacing | Default spacing of pivots within the gap, with denser placement for short gaps | Controls how densely Phase C places pivots within uncovered gaps. |

SSF denotes *singleton suppression filter*; SSF+A denotes *singleton suppression filter with abundance*; CF denotes the suffix array compression factor.

### 3.10. ERT2: Applying supplementary seeding to ERT

During the development of RosaSeed, we realized that the fixed pivot supplementary seeding strategy of Phase B was not specific to RosaSeed and could also be applied to other FM index-based aligners. To investigate this idea, we constructed a simplified ERT variant, which we call **ERT2**, and included it as another baseline alignment algorithm in this study.

The original ERT aligner performs seeding in three stages. ERT2 modifies this pipeline in two ways. First, the original ERT Phase 2 reseeding stage is removed. Second, the original Phase 3 seeding stage is replaced with the supplementary Phase B fixed pivot seeding strategy described in Section 3.5. Specifically, ERT2 evaluates a small number of evenly spaced pivot positions across the read and performs supplementary seeding using the same SA interval stopping criterion employed by Phase B of RosaSeed. Apart from these modifications, ERT2 retains the original ERT index structures and the downstream alignment pipeline remains unchanged. In particular, the radix tree index and associated memory footprint remain identical to those of the original ERT.

### 3.11. Coroutine-Based Prefetch Scheduling

Every FM index backward extension step reads one, and possibly two, occurrence checkpoint blocks at an address(es) determined by ℓ and *h* for the current SA interval [*l, h*). Because successive interval endpoints depend on the symbols matched so far, these addresses ℓ and *h* are not predictable in advance, and the processor’s hardware prefetcher will be ineffective. Each resulting cache miss therefore stalls the processor while a cache line is fetched from main memory. For RosaSeed, which operates on a 49 GB derived reference FM index, this memory latency accounts for the majority of per-read compute time across all the seeding phases.

The new minibwa aligner [16] addresses this potential bottleneck by batching multiple reads together and processing them in a shared pipeline. Rather than completing one read’s SMEM search before beginning the next, minibwa organises extension operations from all reads in the batch into a queue and processes them together in rounds. After each backward extension step, minibwa issues a data prefetch directive for the occurrence array block that the same read’s next extension step will require later, then moves on to the extension step of the next read in the queue. By the time it returns to processing a given read, the prefetched data has had sufficient time to arrive from main memory. minibwa applies the same batching strategy to its SA lookup stage. Taken together, these two optimizations, together with precomputed lookup tables for short *k*-mers, account for minibwa’s reported speedup over BWA-MEM2.

In RosaSeed, we adopted the same latency-hiding prefetching strategy and extended it to all seeding phases and the SA lookup stage. Like minibwa, the scheduler processes a batch of reads together, executing one *s*-base extension per read per round and issuing a data prefetch for the next block before moving to the next read. However, the organization differs: rather than a single unified queue that interleaves operations from all reads across all stages, RosaSeed uses a fixed-size slot array with a round robin iterator, one per seeding phase. This separates the scheduling logic for each phase and allows each phase’s prefetch strategy to target the data structure it needs to access. The following paragraphs describe the scheduler for each phase.

#### Phase A: Cross-read prefetch scheduling

Phase A is RosaSeed’s primary seeding phase. It considers a series of pivots along the *pat*, initialises an SA interval at each pivot using the precomputed JT, and then extends the seed leftward one two-base symbol at a time until the extension fails (the SA interval goes empty). Both the JT lookup and the *s*-base extension steps access the derived reference FM index at addresses that are determined by the current state of the search and cannot be predicted in advance.

To hide the latency of these accesses, we restructured Phase A so that its inner extension loop can be *suspended* after each *s*-base and *resumed* later from exactly the same point. Suspending requires capturing the complete execution state, including the current pivot position, BWT interval endpoints, seed boundaries, selected query strand, and phase indicator, in a per-read state record.

The scheduler processes together a fixed-size batch of reads simultaneously. In each scheduling round it visits every active slot in order, executing exactly one *s*-base extension step per slot. After each step it computes the addresses of the occurrence checkpoint blocks that this slot’s next extension step will require and issues software prefetches for them, then moves to the next slot. After visiting all slots in the batch, it returns to the first. By the time it revisits a given slot, the prefetched blocks have had the duration of all intervening steps to arrive from main memory into the cache.

#### Phase B: within-read and cross-read prefetch scheduling

Phase B places a small number of supplementary pivots at evenly-spaced positions along the read and attempts to find additional seeds from each pivot using the same JT lookup and *s*-base extension steps. Because Phase B operates on only a small number of pivots per read (three by default), the natural unit of parallelism within Phase B was found to be the set of pivots for a single read rather than a set of reads.

The Phase B coroutine therefore interleaves the *s*-base steps *across pivots within one read* : a per-pivot state record captures the extension progress for each pivot, and the scheduler cycles through all active pivots one *s*-base step at a time, issuing prefetches for the next occurrence checkpoint blocks before yielding to the next pivot.

In addition, the function that calls Phase B applies a *cross-read* prefetch: before beginning Phase B for read *i*, function computes the expected first pivot position of reads *i*+1 through *i*+3 and issues prefetches for their first occurrence checkpoint blocks. By the time read *i*’s Phase B completes, the prefetched blocks for the next three reads have had the entire duration of read *i*’s Phase B to arrive from main memory into the cache.

#### Phase C: within-gap prefetch scheduling

Phase C identifies uncovered gaps in the Phase A seed set, that is, the regions of the read where no seed was found. It then attempts to find additional seeds for each gap using gap-directed pivots placed within the gap boundaries. For each gap, Phase C runs one pass on the chosen query direction (that is, either the forward or reverse complement read) and optionally a second pass on the opposite read strand if the corresponding compile-time directive is enabled.

Within each pass, Phase C places multiple pivots within the gap region and applies the same JT and *s*-base seed extension steps as Phases A and B. The number of pivots per gap depends on the gap size and a configurable step parameter. The coroutine for Phase C interleaves *s*-base steps *across pivots within one gap*, using the same per-pivot state record and round-robin structure as Phase B’s within-read scheduler. Data prefetches are issued for the next occurrence checkpoint blocks after each step, hiding FM index access latency within the gap dedicated search.

Phases B and C are mutually exclusive configurations: the codebase supports either fixed-pivot supplementary seeding (Phase B) or gap-directed seeding (Phase C), selected at compile time. The configuration evaluated in this work uses Phase A followed by Phase C, based on the results of extensive alignment experiments.

#### miniRosaSeed: a speed-optimized variant

With all three coroutine schedulers, we also propose *miniRosaSeed* as a more aggressive and faster version of *RosaSeed*. miniRosaSeed reduces the Phase A SA interval cap from *T* = 2000 to *T* = 50, suppressing highly abundant seeds earlier and forwarding fewer candidates to the pre-chain filter, chaining, and BSW extension stages. It also lowers the threshold at which the pre-chain seed suppression heuristic activates. All algorithmic components, the three coroutine schedulers and the BWA-MEM2-compatible downstream pipeline, are identical to the RosaSeed configuration. miniRosaSeed is therefore a point on RosaSeed’s speed-accuracy operating curve rather than a distinct algorithm, and is evaluated below alongside minibwa and RosaSeed in Section 4.7.

### 3.12. Datasets and benchmarking setup

#### Reference genome

All experiments reported below use the Telomere-to-Telomere complete human reference genome assembly T2T-CHM13v2.0 [5]. The assembly consists of 24 chromosomes (chromosomes 1–22, X, and Y), with a total length of approximately 3.1 Gbp, that provides a near-complete representation of the human genome, including highly repetitive regions that are incompletely represented in earlier reference assemblies, such as GRCh38 [32]. The T2T-CHM13v2.0 assembly was chosen because these repetitive regions frequently generate large SA intervals and therefore provide a more demanding benchmark for evaluating the effectiveness of seeding algorithms.

#### Simulated dataset

We also used the ART read simulator [33] to generate a simulated short read dataset at 5× coverage^1^ of T2T-CHM13v2.0, producing approximately 103 million single-ended reads of length 150 bp using ART’s built-in Illumina HiSeq X Ten error profile [33]. The simulator provides ground truth alignment positions for each read, enabling absolute accuracy evaluation (Section 4.4). We will refer to this dataset as the *ART dataset*.

#### Real dataset

We additionally benchmarked all aligners using a real sequencing dataset (accession ERR16657779), downloadable from the European Nucleotide Archive (ENA) [34]. From this accession we extracted 20 million reads and applied the following simplifying filters: reads were trimmed to exactly 150 bp, and any read containing one or more ambiguous bases (so-called *N* bases) was excluded. The filtered dataset thus consists entirely of 150 bp reads over the fourbase alphabet {*A, C, G, T*}. Because true alignment positions are unavailable for real sequencing data, alignment accuracy is evaluated relative to BWA-MEM2 as the reference aligner (Section 4.2). We will refer to this filtered dataset as the *ERR dataset*.

#### HG002 Illumina validation dataset

To assess the downstream variant-calling consistency on a well-characterized benchmark sample, we used sequencing data from HG002 (NA24385), the son of the Ashkenazim Trio and a Genome in a Bottle (GIAB) reference sample [35, 36]. A subset comprising 7,989,074 single-ended Illumina reads (250 bp) was aligned against the complete T2T-CHM13v2.0 reference genome. This subset was selected to provide a consistent evaluation framework for comparing the downstream performance of BWA-MEM2/ERT, minibwa, Strobealign, RosaSeed and miniRosaSeed under identical conditions. We will refer to this dataset as the *HG002 dataset*.

#### Baseline aligners

RosaSeed’s performance was compared against the following aligners, each run using its recommended settings for short Illumina reads:

- **BWA-MEM2 (uncompressed SA)**: BWA-MEM2 using an uncompressed suffix array index. This configuration serves as the baseline aligner for speed comparisons and the reference aligner for accuracy evaluation.
- **BWA-MEM2 (default compressed SA)**: BWA-MEM2 using its default compressed SA with a compression factor of 8 [11].
- **ERT** [10]: The Enumerated Radix Tree aligner using its standard 15-mer prefix index.
- **ERT2**: Simplified ERT-derived baseline introduced in Section 3.10 477 cl:32.
- **Minimap2** [8]: Run using the short read preset `-ax sr -k19` `-w10`.
- **Bowtie2** [6]: Run using default parameters.
- **Strobealign** [13]: Run using default parameters with precomputed index.
- **minibwa** [16]: Run using default parameters.

##### Hardware and threading

The experiments reported in Sections 4.1–4.6 were conducted on a Lenovo ThinkStation P620 equipped with an AMD Ryzen Threadripper PRO 3945WX processor (12 physical Zen 2 cores, 24 hardware threads, 6-MB L2 Cache, 64-MB L3 cache) and approximately 300 GB of DDR4-3200 ECC memory running Ubuntu 20.04. The later experiments reported in Sections 4.7 and 4.8 were conducted on an upgraded Lenovo ThinkStation P620 equipped with an AMD Ryzen Threadripper PRO 5965WX processor (24 physical Zen 3 cores, 48 hardware threads, 12-MB L2 Cache, 128-MB L3 cache) and approximately 300 GB of DDR4 memory running Ubuntu 26.04.

The per-read seeding time and memory footprint are reported from single-threaded runs to isolate algorithmic cost and eliminate parallelism effects. Single-threaded execution times were measured using wall clock time and averaged across five independent runs. While a warm cache was maintained before each run, the execution lengths ensured that initial cache states did not meaningfully skew the overall averages. The memory footprint was recorded as the peak resident set size (RSS) reported by the operating system during the single-threaded run. The fastq-to-SAM runtime (time measured from when the aligner starts to the time when it finishes writing the output SAM file) is additionally reported for increasing thread counts ∈ {1, 2, 4, 6, 8, 10, 12, 14, 16, 18, 20, 22, 24} to characterize multi-threaded scaling behavior in section 4.5. All of the aligners were run under identical hardware and OS conditions with the same thread count to ensure a fair comparison.

### 3.13. Alignment Accuracy Metrics

We evaluated alignment accuracy using three metrics, each targeting a different aspect of alignment quality. For all three metrics, only primary alignments (as so designated by the BWA-MEM2 default pipeline parameters) are considered; secondary and supplementary alignments are excluded from evaluation. For consistency across nonstandard metrics, reads for which the reference baseline is unmapped (BWA-MEM2) are excluded from structural and sequence-consistent accuracy computation. The three considered definitions of alignment accuracy are given below.

*Standard accuracy* measures whether the reported alignment falls within 50 bp of the true position in the reference, following the convention established by Bowtie2 [6], and is our primary accuracy measure. A read is considered correctly mapped if the aligner and truth alignments agree on the read strand and reference sequence, and their leftmost start positions satisfy | start(*a*^⋆^) − start(*a*) | ≤ 50, where *a*^⋆^ denotes the truth alignment and *a* is the aligner alignment. We also report the *unmapped read fraction*, defined as the fraction of reads for which the aligner reported the read as unmapped. Standard accuracy has an important limitation: it treats all incorrect placements identically. For example, a read mapped to a different chromosome and a read mapped 60 bp away within a tandem repeat region are both counted as errors, even though the latter may correspond to a plausible placement within a repetitive region of the reference.

*Structural accuracy* addresses a limitation of standard accuracy by evaluating whether the aligner produces an alignment with a comparable edit burden to the ground truth, regardless of the reported genomic reference positions. For an alignment *r*, the *edit burden* is defined to be *E*(*r*) = *I*(*r*) + *D*(*r*) + *X*(*r*), where *I*(*r*), *D*(*r*), and *X*(*r*) are the number of inserted bases, deleted bases, and explicit mismatch operations, respectively, as recorded in the CIGAR string [23]. An alignment is *structurally correct* if | *E*(*a*^⋆^) − *E*(*a*) | */ N* ≤ 0.05, where *N* is the read length. The 5% threshold permits modest differences in local alignment representation while still requiring close agreement between the reported and groundtruth edit burdens. This metric is evaluated only over reads for which the aligner produced a mapped primary alignment. Two alignments mapping to different copies of a repeat with identical edit burden are considered to be structurally equivalent and thus counted as correct, thus distinguishing genuine misalignment from equally plausible placements in repetitive regions.

*Sequence-consistent accuracy* (SCA) measures whether the reported alignment is biologically plausible by evaluating the agreement between the read and the reference at the reported alignment location. Specifically, SCA checks whether the normalized edit distance between the read and the reference at the reported position satisfies NM(*a*)*/N* ≤ 0.10, where NM(*a*) is obtained from the SAM `NM` tag (NM: edit distance to the reference). When the `NM` tag is absent, the edit distance is approximated as *I*(*a*)+*D*(*a*)+*X*(*a*) from the CIGAR string. For a 150-bp read, this threshold permits up to 15 edits, substantially exceeding the expected sequencing error rate of modern Illumina data [20]. Thus SCA captures a failure mode that index-based metrics can miss: a high-edit distance placement within 50 bp of the true alignment is still a relatively poor quality result. SCA gives the fraction of mapped primary reads satisfying this criterion.

For the ART-simulated read dataset, the simulator-provided reference positions served as the ground truth *a*^⋆^ for all three metrics. For the ERR real dataset, where true reference positions are unavailable, BWA-MEM2 primary alignments serve as the ground truth. It is to be noted that BWA-MEM2 therefore, by definition, achieves perfect standard accuracy. Accuracy results on real data should therefore be interpreted as measuring the alignment consistency with respect to BWA-MEM2 (the widely used standard aligner) rather than absolute correctness. Together, the three metrics assess complementary aspects of alignment quality. Standard accuracy evaluates positional correctness relative to the ground truth; structural accuracy evaluates agreement between the reported and ground-truth alignment edit structures; and sequence-consistent accuracy evaluates the biological plausibility of the reported alignment by requiring a low edit distance between the read and the reference sequence at the reported genome location.

## 4. Results

### 4.1. Experimental Overview

The RosaSeed framework provides a multidimensional design space through configurable symbol multiplicity (the number of bases per extension step), supplementary seeding strategy (Phase B or C), SA compression, gap-directed policy (Phase C), and pre-chain filtering heuristics. Rather than presenting the full design space for all possible parameter combinations, our experimental evaluation will focus on four representative configurations selected to illustrate the principal operating regimes of the RosaSeed framework.

*RosaSeed* will refer to the recommended operating configuration. This configuration provides a strong overall balance between runtime, memory footprint, and alignment accuracy. *RosaSeed-FixedPivots* employs the fixed-position supplementary seeding strategy introduced in Phase B and serves as a representative fixed-pivot design. *RosaSeed-3Base* demonstrates the effect of coarsegrained three-base FM index traversal on seeding efficiency, whereas *RosaSeed-Compact* emphasizes reduced memory consumption through aggressive suffix array compression while maintaining competitive alignment performance.

The following subsections first compare these four representative operating configurations against established short read aligners before examining the contribution of individual algorithmic components through controlled ablation studies.

### 4.2. Overall Benchmark Comparison

Figure 7 and Table 4 compare the four representative RosaSeed configurations against widely used short read aligners on the ERR dataset. Taken together, the results show that the RosaSeed framework supports multiple operating regimes, covering both performance-oriented and memory-efficient versions, while consistently delivering high alignment quality.

**Table 4.** Runtime, memory footprint, unmapped read fraction, and alignment accuracy for the ERR dataset. Runtime measurements are reported in microseconds per read (*µ*s/read) for single-threaded execution on an AMD 12-core Zen 2 workstation. The *seed-to-BSW* time denotes the combined execution time of seed generation, suffix array coordinate retrieval, chaining and Banded Smith–Waterman (BSW) extension. Seed processing time denotes the profiled time associated with seed generation and reference position retrieval/sorting stages, where available. For aligners whose implementations do not expose these stages separately, the closest corresponding profiled component is reported.

| Configuration | fastq-to-SAM<br>runtime<br>($\mu\text{s}/\text{read}$) | seed-to-BSW<br>runtime<br>($\mu\text{s}/\text{read}$) | Seed-proc.<br>time<br>($\mu\text{s}/\text{read}$) | Memory<br>footprint<br>(GB) | Unmapped<br>reads<br>(%) | Std.<br>acc.<br>(%) | Struct.<br>acc.<br>(%) | SCA<br>(%) |
| --- | --- | --- | --- | --- | --- | --- | --- | --- |
| *Baseline aligners* |  |  |  |  |  |  |  |  |
| Bowtie2 | 378.78 | 372.24 | 172.18 | 4.20 | 0.803 | 93.81 | 99.93 | 99.71 |
| Minimap2 | 115.61 | 104.17 | 24.99 | 12.89 | 0.803 | 93.94 | 99.90 | 99.83 |
| BWA-MEM2 (SA cf=8) | 141.22 | 135.61 | 77.82 | 16.20 | 0.374 | 100.00 | 100.00 | 99.88 |
| BWA-MEM2 (uncompr. SA) | 134.79 | 127.12 | 67.63 | 41.70 | 0.374 | 100.00 | 100.00 | 99.88 |
| ERT | 86.57 | 79.11 | 20.50 | 66.30 | 0.374 | 100.00 | 100.00 | 99.88 |
| ERT2 | 50.26 | 44.11 | 6.20 | 66.30 | 0.374 | 99.63 | 99.99 | 99.88 |
| *Representative RosaSeed configurations* |  |  |  |  |  |  |  |  |
| RosaSeed | **34.97** | 29.98 | 5.18 | 49.61 | 0.523 | 98.52 | 99.97 | 99.87 |
| RosaSeed-3Base | 41.12 | **28.81** | **3.75** | 142.62 | 0.523 | 98.52 | 99.97 | 99.87 |
| RosaSeed-FixedPivots | 43.93 | 38.58 | 7.51 | 49.63 | 0.552 | 98.54 | **99.98** | **99.88** |
| RosaSeed-Compact | 44.34 | 39.90 | 9.15 | **25.86** | **0.382** | **98.61** | **99.98** | **99.88** |

Bold values indicate the best result among the representative RosaSeed configurations for the corresponding metric.

**Figure 7.**
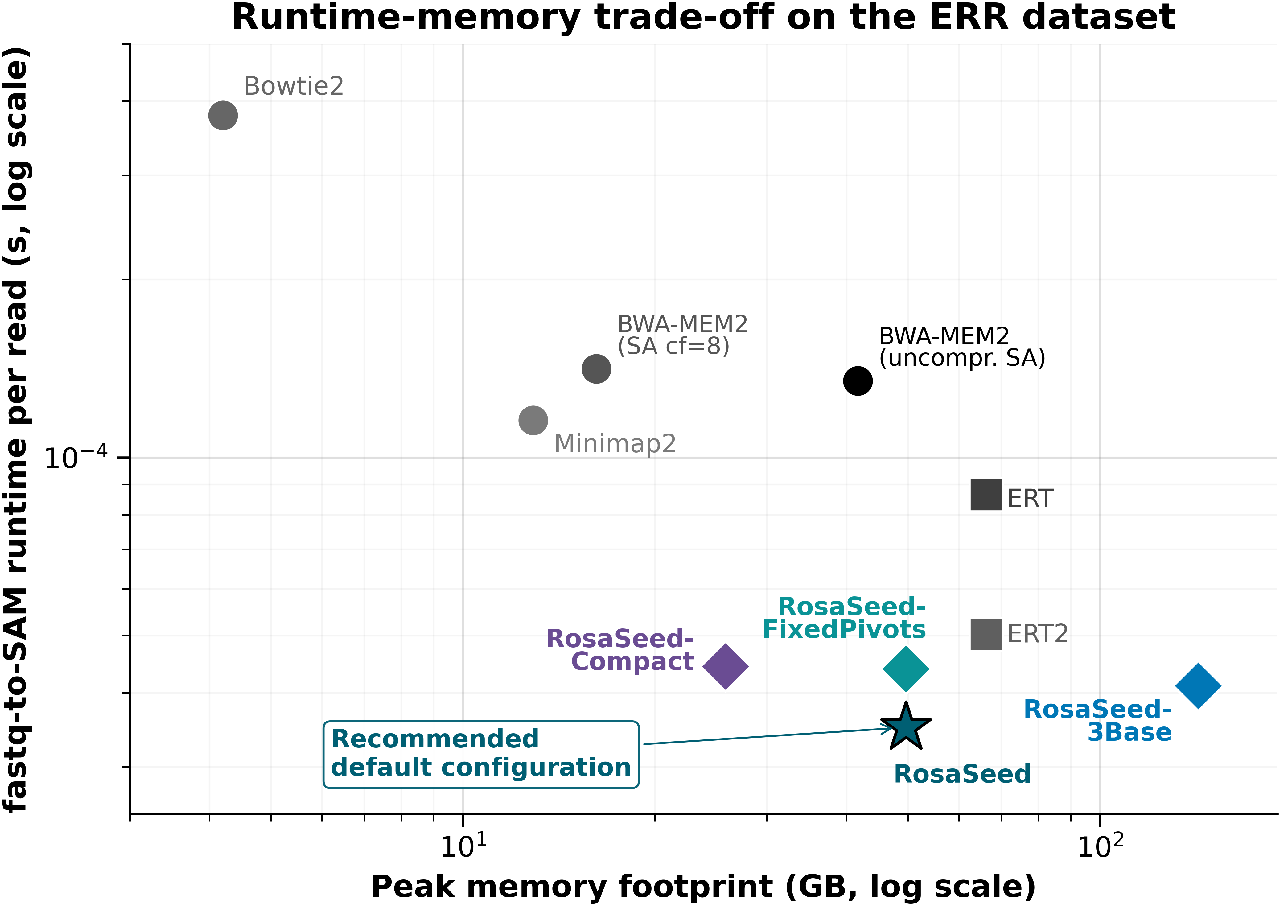
Runtime-memory trade-off of the four representative RosaSeed configurations compared with established short read aligners on the ERR dataset (20 million reads, 150 bp, single-threaded execution). The horizontal axis shows the peak memory footprint, while the vertical axis reports the fastq-to-SAM average alignment runtime per read. Smaller memory footprint and shorter average runtime per read correspond to better performance. RosaSeed is highlighted as the recommended operating configuration, providing the best overall balance between runtime, memory footprint, and alignment accuracy among the four representative RosaSeed operating modes.

#### RosaSeed (recommended configuration)

Among the representative configurations, RosaSeed achieves the most favorable overall balance between runtime, memory footprint, and alignment accuracy and is therefore selected as the recommended operating configuration. As shown in Figure 7, RosaSeed sits on the runtime-memory Pareto frontier among the representative RosaSeed configurations, achieving the lowest fastq-to-SAM runtime while requiring substantially less memory than either ERT or ERT2.

Using fastq-to-SAM runtime (assuming single-threaded execution) as the primary performance metric, RosaSeed completes alignment at an average rate of only 34.97 *µ*s/read, corresponding to speedups of 2.48× over ERT, 1.44× over ERT2, 3.31× over Minimap2, 4.04× over BWA-MEM2 (SA cf=8), 3.85× over BWA-MEM2 with an uncompressed SA, and more than 10.83× over Bowtie2. Despite these runtime improvements, RosaSeed requires only 49.61 GB of peak memory, which is roughly 25% less than both ERT and ERT2.

The improvement in fastq-to-SAM runtime is largely driven by a more efficient seeding strategy. RosaSeed requires only 5.18 *µ*s/read for seed processing, making it approximately 3.96×, 1.20×, 13.06×, 15.02×, 4.82×, and 33.24× faster at seed processing than ERT, ERT2, BWA-MEM2 with an uncompressed SA, BWA-MEM2 (SA cf=8), Minimap2 and Bowtie2, respectively. The seed-to-BSW (time required for seed processing + SAL + chaining + BSW) rate is also reduced to 29.98 *µ*s/read, showing that the performance gains extend beyond seed generation to the short read alignment computation as a whole. Importantly, these computational improvements come with negligible loss of alignment accuracy. RosaSeed maintains 98.52% standard accuracy, 99.97% structural accuracy, and 99.87% sequence-consistency accuracy, remaining closely competitive with the most accurate benchmark aligners while providing substantially faster alignment speed. These results together establish RosaSeed as a significant advance over all the contemporary alignment algorithms. RosaSeed uses a Phase A SA-interval cap of 2000 for this recommended configuration; however, a cap of 5000 can be used to reduce the unmapped rate from 0.523% to 0.385%, at the cost of increased alignment time.

#### RosaSeed-3Base

RosaSeed-3Base demonstrates the impact of 3-base symbol traversal on seeding performance. Because each traversal step extension processes a larger multiplicity alphabet with more bases considered in each step, RosaSeed-3Base significantly reduces the time required during backward search and therefore represents the performance-oriented extreme of the RosaSeed design space.

Across all representative configurations, RosaSeed-3Base records the fastest seed processing time at 3.75 *µ*s/read and the lowest seedto-BSW time at 28.81 *µ*s/read, outperforming even the recommended RosaSeed configuration on both metrics. Compared to BWA-MEM2 with an uncompressed SA, seed processing is roughly 18.0× faster. It also achieves speedups of 5.47× over ERT, 1.65× over ERT2, 6.66× over Minimap2, and 45.9× over Bowtie2.

Although RosaSeed-3Base reports the fastest seed processing and seed-to-BSW execution among all representative configurations, the larger FM index required for the derived 3-base reference increases the memory footprint to 142.62 GB. As a result, the fastq-to-SAM runtime advantage that RosaSeed-3Base holds over RosaSeed is partially lost by the greater index footprint and data movement costs. These findings show that while increasing the symbol multiplicity can substantially speed up seeding and the seed-to-BSW kernel, it makes the fastq-to-SAM alignment processing slower as there is a cost to load a larger memory footprint for the FM index.

#### RosaSeed-FixedPivots

RosaSeed-FixedPivots implements the fixed-position supplementary seeding strategy of Phase B, offering an alternative operating point within the RosaSeed framework. While employing a simpler supplementary seeding mechanism than RosaSeed, this configuration maintains competitive runtime and alignment accuracy, thereby illustrating the effectiveness of fixed-pivot supplementary seed generation.

RosaSeed-FixedPivots achieves a fastq-to-SAM alignment runtime of 43.93 *µ*s/read, corresponding to speedups of 1.97× over ERT, 1.14× over ERT2, 2.63× over Minimap2, 3.21× over BWA-MEM2 (SA cf=8), 3.07× over BWA-MEM2 with an uncompressed SA, and roughly 8.62× over Bowtie2. Its peak memory footprint of 49.63 GB remains substantially smaller than that of ERT and ERT2 while requiring essentially the same memory space as RosaSeed. RosaSeed-FixedPivots requires 7.51 *µ*s/read for seed processing and 38.58 *µ*s/read for the seed-to-BSW kernel, both of which remain well below the corresponding figures for the benchmark aligners.

With respect to accuracy, RosaSeed-FixedPivots (Phase A & B) achieves 98.54% standard accuracy, 99.98% structural accuracy, and 99.88% sequence-consistency accuracy. These results confirm that fixed-position supplementary seeding is a competitive option within the RosaSeed framework, although the adaptive gap-directed strategy used in recommended RosaSeed (Phase A & C) achieves the same accuracy and is 20.4% faster.

#### RosaSeed-Compact

RosaSeed-Compact is an especially memory-efficient configuration within the RosaSeed framework, pairing conventional 1-base FM index traversal with aggressive SA compression by a factor of 8. It is intended for computing environments (say, an older workstation) where memory capacity is the main constraint, without giving up competitive alignment performance. RosaSeed-Compact requires only 25.86 GB of peak memory, the smallest footprint among the representative RosaSeed configurations. RosaSeed-Compact reduces the memory footprint by roughly 48% compared to RosaSeed, 61% compared to ERT/ERT2, and 82% compared to RosaSeed-3Base. Despite this significant drop in memory footprint, it has an alignment time of just 44.34 *µ*s/read, making it 1.95× faster than ERT, 1.13× faster than ERT2, 2.61× faster than Minimap2, 3.18× faster than BWA-MEM2 (SA cf=8), and more than 8.54× faster than Bowtie2. The reduced index size causes a modest increase in seed processing time to 9.15 *µ*s/read relative to the higher-performance RosaSeed configurations, reflecting the trade-off between compression and search efficiency. Nevertheless, RosaSeed-Compact still maintains 98.61% standard accuracy, 99.98% structural accuracy, and 99.88% sequence-consistency accuracy. This shows that meaningful memory savings are achievable at only a modest runtime cost relative to the higher-performance RosaSeed configurations.

### 4.3. Design Evaluation and Ablation Study

#### Effect of adaptive supplementary seeding

To assess how much supplementary seed generation contributes to alignment speed and accuracy, three seeding strategies are compared. The first uses the fixed-pivot supplementary seeding approach from Phase B, which evaluates a small set of predetermined pivot positions. The second is the recommended RosaSeed implementation used throughout this work, which applies gap-directed exploration in both alignment directions and includes early termination once the current gap is covered. The third is an adaptive gap-directed strategy that checks the reverse alignment direction only when extra seed recovery is needed. Together, these configurations represent a progression from fixed supplementary pivots toward increasingly thorough gap-directed seed generation, and they allow the associated performance trade-offs to be measured directly.

Table 5 summarizes how each strategy affects the seed-to-BSW time, seed processing time, and memory footprint. Moving from (1) the fixed-pivot strategy of Phase B to (2) the adaptive gap-directed supplementary seeding brings the seed-to-BSW time down from 38.58 *µ*s/read to 29.98 *µ*s/read and the seed processing time from 7.51 *µ*s/read to 5.18 *µ*s/read, with virtually no change in memory usage. Rather than checking a fixed set of supplementary pivot locations, the adaptive strategy directs supplementary search effort only toward uncovered regions identified after the primary seeding stage (Phase A), focusing the computation where new seeds are most likely to improve the alignment quality. This targeted approach reduces the seed processing overhead and also lowers the overall seed-to-BSW time by cutting down the number of candidate alignments that need downstream processing, showing that better seed placement provides benefits in the later alignment stages.

**Table 5.** Comparison of three supplementary seeding strategies. The default adaptive gap-directed strategy (2) achieves the lowest runtime while preserving essentially the same memory footprint, demonstrating that effective gap-directed supplementary seed generation can be obtained with relatively small additional computational overhead.

| Strategy | seed-to-BSW<br>runtime<br>($\mu\text{s}/\text{read}$) | Seed<br>processing<br>($\mu\text{s}/\text{read}$) | Memory<br>(GB) |
| --- | --- | --- | --- |
| (1) Phase B (fixed pivots) | 38.58 | 7.51 | 49.63 |
| (2) Adaptive gap-directed (**RosaSeed**) | **29.98** | **5.18** | 49.59 |
| (3) Adaptive gap-directed (conditional opposite direction) | 30.42 | 5.24 | 49.61 |

Note: Bold values indicate the best runtime result for the corresponding metric.

The two adaptive gap-directed variants show remarkably similar performance. The RosaSeed configuration, strategy (2) in Table 5, which evaluates both alignment directions and applies early termination, records the lowest seed-to-BSW time (29.98 *µ*s/read) and seed processing time (5.18 *µ*s/read), marginally outperforming the conditional opposite-alignment-direction strategy (3) despite performing a more thorough supplementary search. This indicates that the additional cost of the strategy is effectively amortized by the early exit mechanism, allowing more complete supplementary seed recovery without introducing significant computational overhead.

##### Effect of s-base symbol multiplicity

To evaluate the impact of multi-base alignment, three configurations employing one-, two- and three-base traversal are compared while keeping all other algorithmic parameters identical. *RosaSeed* is used with a 15-base JT, and its equivalent one-base and 3-base configurations are compared. Specifically, every configuration uses the same 15-base JT, a SA compressed by a factor of 2, the Phase C gap-directed supplementary seeding strategy and pre-chain seed filtering heuristics, so any observed differences can be attributed solely to the use of multi-base symbols.

Figure 8 presents the seed-to-BSW time, seed processing time, and memory footprint produced by each of the three traversal strategies. As the degree of multi-base alignment increases from 1-base to 3-base, both the seed processing stage and the overall seed-to-BSW become progressively faster. Seed processing time drops from 6.12 *µ*s/read with a 1-base sweep to 4.89 *µ*s/read with a 2-base sweep and then further down to 4.33 *µ*s/read with a 3-base sweep, representing a time reduction of roughly 29% between the 1-base and 3-base configurations. The seed-to-BSW time follows the same trend, falling from 31.07 *µ*s/read to 30.42 *µ*s/read and then to 29.75 *µ*s/read as multiplicity increases from one base per step to three.

**Figure 8.**
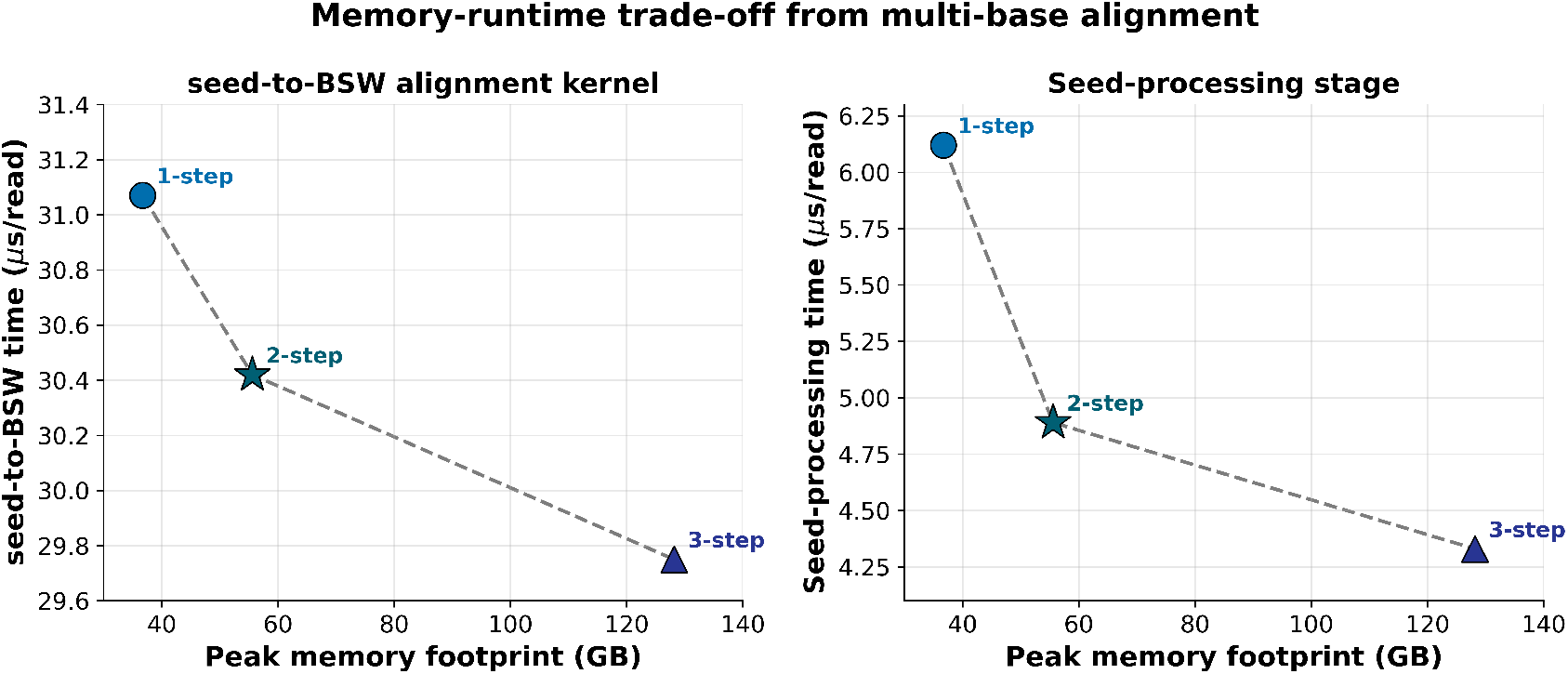
Effect of multi-base alignment on runtime (single thread) and memory footprint. The three configurations employ 1-base, 2-base, and 3-base sweeps while keeping all other algorithmic parameters identical. The left plot reports the seed-to-BSW time (comprising seed generation, suffix array retrieval, and BSW extension), whereas the right plot reports the seed processing time. Increasing the symbol multiplicity progressively reduces both seed processing and seed-to-BSW time at the expense of a larger memory footprint, illustrating the inherent memory-performance trade-off of multi-base alignment. The 2-base configuration adopted by recommended RosaSeed can be seen to provide an effective compromise between runtime and memory consumption.

These gains occur because greater-multiplicity symbol representations pack multiple bases into a single FM index sweep step, which reduces the number of backward search iterations needed during seed generation. Fewer FM index lookups are therefore required to build candidate seeds, lowering seed processing overhead and reducing the execution time in the downstream phases.

However, the runtime improvements come at the cost of a considerably larger memory footprint. Peak memory usage rises from 36.74 GB for the 1-base configuration to 55.58 GB for the 2-base configuration, and to 128.16 GB for the 3-base configuration, driven by the larger derived-reference FM index that multiple-base symbol encodings require.

Although the 3-base configuration achieves the shortest seed processing and seed-to-BSW times, its memory footprint is three times larger than that of the 1-base configuration. The 2-base design, by contrast, captures most of the available runtime benefit while consuming substantially less memory than the 3-base design. For this reason, the 2-base traversal adopted in recommended RosaSeed offers a favorable balance between computational efficiency and memory use.

##### Effect of suffix array compression

To assess how SA compression affects performance, four configurations are compared using compression factors of 1, 2, 4 and 8, with all other algorithmic parameters held constant. A compression factor of 1 means the SA is stored without compression, while higher factors progressively reduce the number of stored samples and instead reconstruct SA values (reference positions) as needed during alignment. As a result, increasing the compression factor is expected to lower the memory usage at the cost of additional computation for reconstructing reference positions.

Figure 9 presents the resulting trade-offs across the four compression factors, covering memory footprint, seed-to-BSW time, and seed processing time. Note how SA compression involves a trade-off between memory usage and runtime. The SA memory footprint drops from 29.04 GB with the exact suffix array (compression factor 1) to 14.52 GB, 7.26 GB, and 3.63 GB for compression factors of 2, 4 and 8, respectively. As a result, the total memory footprint of RosaSeed drops from 64.13 GB with the exact suffix array (compression factor 1) to 49.61 GB, 42.35 GB and 38.72 GB for SA compression factors of 2, 4 and 8, respectively. This amounts to a total memory reduction of nearly 40% compared to the uncompressed representation.

**Figure 9.**
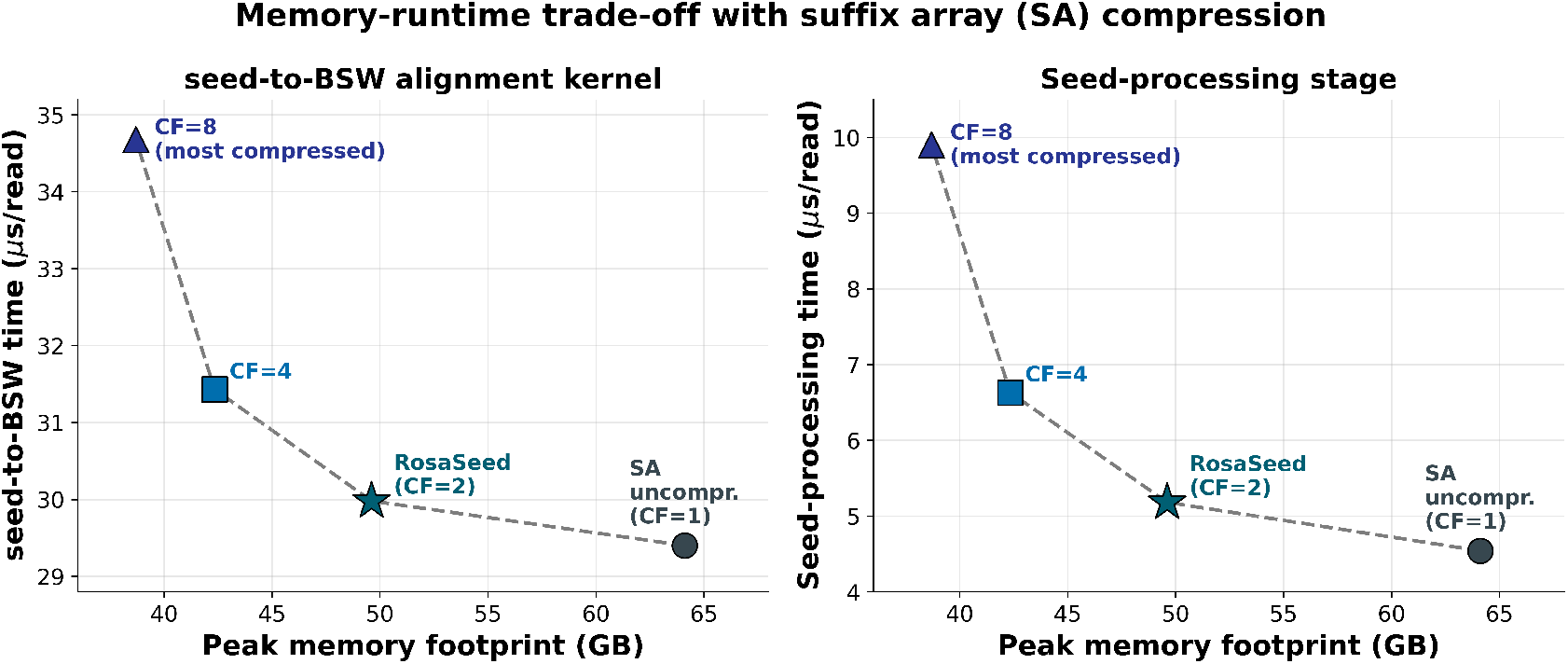
Memory and runtime (single thread) trade-offs for suffix array compression. The four configurations use suffix array compression factors of 1 (uncompressed suffix array), 2, 4 and 8, with all other algorithmic parameters held constant. The left panel shows the seed-to-BSW time and the right panel shows the seed processing time. As the compression factor increases, peak memory usage decreases, but reconstructing SA positions takes longer, resulting in longer runtimes. The compression factor of 2 used in RosaSeed offers a reasonable balance between computational efficiency and memory footprint.

This memory saving comes at the cost of a gradual increase in computation time. The seed-to-BSW time rises from 29.40 *µ*s/read for the exact suffix array to 29.98 *µ*s/read, 31.43 *µ*s/read, and 34.68 *µ*s/read as the compression factor increases from 2 to 8. The seed processing stage shows an even steeper trend, with its runtime growing from 4.54 *µ*s/read for the exact suffix array to 5.18 *µ*s/read, 6.63 *µ*s/read, and 9.90 *µ*s/read for compression factors of 2, 4, and 8, respectively. This pattern is expected since SA compression mainly affects SA lookup operations, where reference positions must be reconstructed before candidate seeds can be passed to the downstream alignment stages. As a result, this reconstruction overhead shows up first in the seed processing time and then feeds into the overall seed-to-BSW runtime operations.

Among all the configurations tested, the compression factor of 2 used by RosaSeed offers the best overall balance. Relative to using an uncompressed SA, the compression factor of 2 cuts the total memory usage by roughly 23% (from 64.13 GB to 49.61 GB) while adding only about 2% to the seed-to-BSW time (from 29.40 *µ*s/read to 29.98 *µ*s/read).

##### Effect of pre-chain seed filtering

To evaluate the contribution of the proposed pre-chain seed filtering heuristics, three different filtering strategies are evaluated: (1) a baseline without pre-chain filtering, (2) a weak singleton suppression filter (SSF), and (3) the abundance-aware variant (SSF+A). SSF suppresses weak singleton-chain candidates once the number of chains already created for a read exceeds a predefined threshold and the candidate seed length falls below a specified minimum. SSF+A builds on this criterion by additionally considering SA interval abundance, suppressing only those short singleton seeds (i.e., seeds that would otherwise initiate chains of length one) whose abundance exceeds a threshold. All other algorithmic parameters remain fixed.

Table 6 summarizes how the proposed pre-chain seed filtering heuristics affect downstream alignment computation. Since filtering is applied only after seed generation and reference position recovery, the seed processing stage itself stays largely unchanged across all tested configurations. The heuristics instead act at the chain construction cap by suppressing low-value singleton seed candidates before they can initiate new chains. The SSF strategy achieves the greatest reduction in downstream work, yielding the lowest seedto-BSW time at 29.98 *µ*s/read. The SSF+A strategy introduces an additional consideration by accounting for seed abundance, retaining short singleton seeds that align at relatively few reference locations while preferentially discarding those that appear to align to highly repetitive regions of the reference.

**Table 6.**
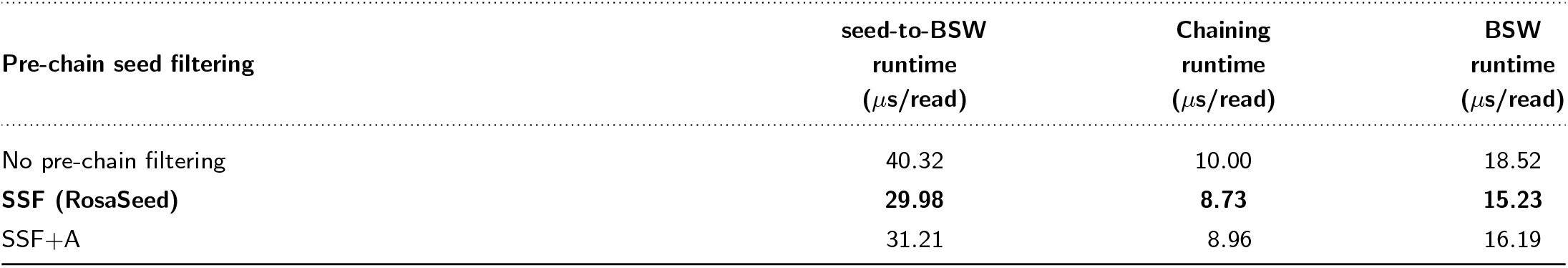
Effect of pre-chain seed filtering on downstream alignment computation time measured for 150-bp short reads in the ERR dataset. Pre-chain filtering reduces chaining and BSW extension work by suppressing low-value, single-seed chain candidates before chain construction.

| Pre-chain seed filtering | seed-to-BSW<br>runtime<br>($\mu\text{s}/\text{read}$) | Chaining<br>runtime<br>($\mu\text{s}/\text{read}$) | BSW<br>runtime<br>($\mu\text{s}/\text{read}$) |
| --- | --- | --- | --- |
| No pre-chain filtering | 40.32 | 10.00 | 18.52 |
| **SSF (RosaSeed)** | **29.98** | **8.73** | **15.23** |
| SSF+A | 31.21 | 8.96 | 16.19 |

### 4.4. Evaluation on Simulated Datasets

To complement the evaluation on real short read data, all representative RosaSeed configurations and baseline aligners were also tested on a simulated short read dataset with known ground truth mapping locations. Unlike the real dataset, the simulated benchmark allows direct measurement of mapping correctness while still providing runtime and memory footprint comparisons.

Table 7 summarizes the fastq-to-SAM alignment runtime, seed-to-BSW time, seed processing time, peak memory footprint, unmapped-read fraction, and alignment accuracy for six baseline aligners and the four RosaSeed configurations. The overall performance trends closely mirror those observed on the real dataset. RosaSeed achieves the shortest fastq-to-SAM runtime among the representative RosaSeed configurations at 28.10 *µ*s/read, using 49.6 GB of memory. RosaSeed-3Base records the shortest seed processing time (3.94 *µ*s/read) and the shortest seed-to-BSW time (24.57 *µ*s/read), which highlights how effective multi-base FM index traversal is for accelerating seed generation. However, this performance gain comes at the cost of a considerably larger memory footprint of 142.82 GB, once again illustrating the runtime-memory trade-off that comes with increased symbol multiplicity. RosaSeed-Compact sits at the opposite end of the design space, requiring only 27.24 GB of memory while still delivering competitive runtime. In contrast, RosaSeed-FixedPivots represents a middle ground based on the simpler fixed supplementary seeding strategy.

**Table 7.**
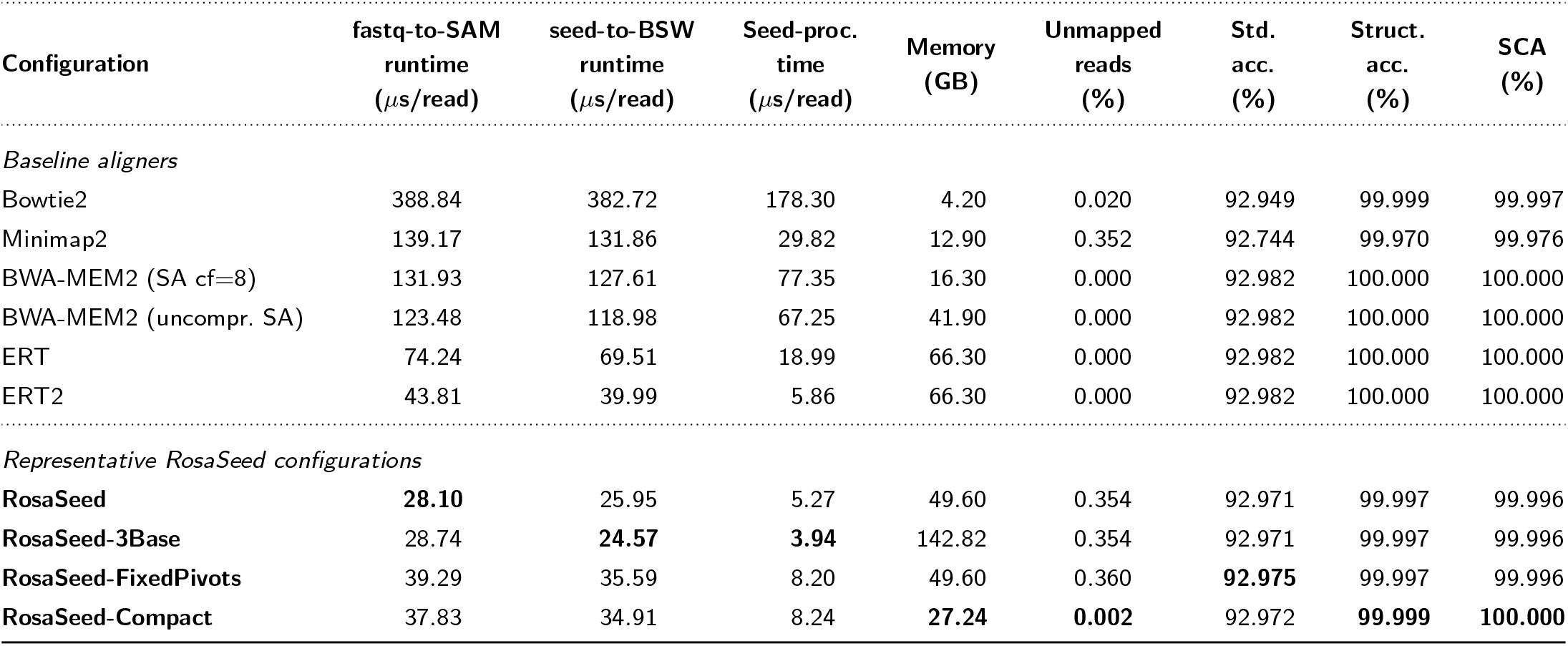
Runtime, memory footprint, fraction of unmapped reads, and alignment accuracy on the simulated ART dataset. Runtime measurements are reported in microseconds per read (*µ*s/read) for single-threaded execution. The *seed-to-BSW* time gives the combined execution time of seed generation, suffix array coordinate retrieval, chaining and Banded Smith–Waterman (BSW) alignment extension. Seed processing time denotes the profiled time associated with seed generation and reference position retrieval/sorting stages, where available. For aligners whose implementations do not provide these stages separately, the closest corresponding profiled component is reported.

| Configuration | fastq-to-SAM<br>runtime<br>($\mu\text{s}/\text{read}$) | seed-to-BSW<br>runtime<br>($\mu\text{s}/\text{read}$) | Seed-proc.<br>time<br>($\mu\text{s}/\text{read}$) | Memory<br>(GB) | Unmapped<br>reads<br>(%) | Std.<br>acc.<br>(%) | Struct.<br>acc.<br>(%) | SCA<br>(%) |
| --- | --- | --- | --- | --- | --- | --- | --- | --- |
| *Baseline aligners* |  |  |  |  |  |  |  |  |
| Bowtie2 | 388.84 | 382.72 | 178.30 | 4.20 | 0.020 | 92.949 | 99.999 | 99.997 |
| Minimap2 | 139.17 | 131.86 | 29.82 | 12.90 | 0.352 | 92.744 | 99.970 | 99.976 |
| BWA-MEM2 (SA cf=8) | 131.93 | 127.61 | 77.35 | 16.30 | 0.000 | 92.982 | 100.000 | 100.000 |
| BWA-MEM2 (uncompr. SA) | 123.48 | 118.98 | 67.25 | 41.90 | 0.000 | 92.982 | 100.000 | 100.000 |
| ERT | 74.24 | 69.51 | 18.99 | 66.30 | 0.000 | 92.982 | 100.000 | 100.000 |
| ERT2 | 43.81 | 39.99 | 5.86 | 66.30 | 0.000 | 92.982 | 100.000 | 100.000 |
| *Representative RosaSeed configurations* |  |  |  |  |  |  |  |  |
| RosaSeed | **28.10** | 25.95 | 5.27 | 49.60 | 0.354 | 92.971 | 99.997 | 99.996 |
| RosaSeed-3Base | 28.74 | **24.57** | **3.94** | 142.82 | 0.354 | 92.971 | 99.997 | 99.996 |
| RosaSeed-FixedPivots | 39.29 | 35.59 | 8.20 | 49.60 | 0.360 | **92.975** | 99.997 | 99.996 |
| RosaSeed-Compact | 37.83 | 34.91 | 8.24 | **27.24** | **0.002** | 92.972 | **99.999** | **100.000** |

The simulated benchmark further confirms that the computational advantages of RosaSeed are achieved without significant degradation in alignment accuracy. All representative RosaSeed configurations achieve structural accuracies of 99.997–99.999% and sequence-consistency agreement of 99.996–100.000%, remaining essentially identical in accuracy to the baseline aligners. Standard alignment accuracy also varies only slightly across RosaSeed configurations (92.971-92.975%) and stays within 0.011 percentage points of the values produced by BWA-MEM2, ERT and ERT2. Although the recommended RosaSeed configuration and RosaSeed-3Base show a modestly higher unmapped-read fraction (0.354%), these reads make up only a small portion of the dataset and therefore have a negligible effect on overall alignment quality, which explains the close agreement across all three accuracy metrics.

The higher unmapped read fraction reflects a deliberate design decision rather than a fundamental limitation of the algorithm. The RosaSeed configuration uses a relatively aggressive Phase A SA interval cap of 2000 to maximize throughput. Raising this cap to 5000 brings the unmapped-read fraction down to just 0.002% with only a modest increase in runtime, showing that alignment accuracy can often be improved through parameter tuning. Taken together, these results indicate that the alternative seeding strategies used by RosaSeed preserve downstream alignment accuracy while offering a tunable runtime-accuracy trade-off that can be adjusted to suit different application requirements.

### 4.5. Scalability of Multithreaded Execution

To assess parallel scalability, all the baseline aligners and the recommended RosaSeed configuration were run to align the ERR dataset on a Lenovo P620 12-core Zen 2 workstation with application thread counts ranging from 1 to 24. Figure 10 shows how the fastq-to-SAM alignment runtime varies with the thread count.

**Figure 10.**
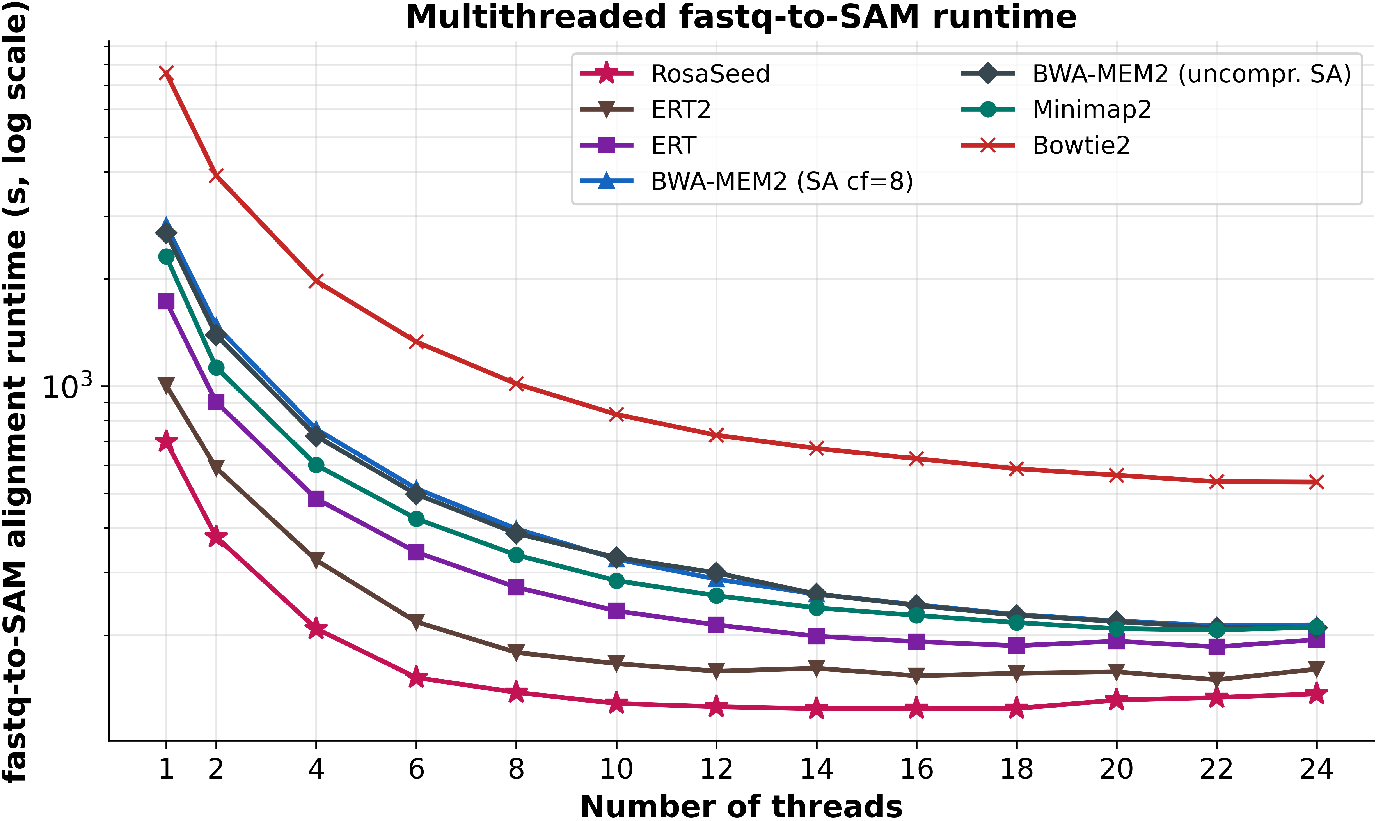
Multithreaded fastq-to-SAM alignment runtime versus number of application threads for baseline aligners and RosaSeed on the ERR dataset. Each point of the plot is the mean of 12 subsequent runs. Runtime is reported on a logarithmic scale. Lower values indicate better performance.

All the tested aligners show substantial runtime reductions as the thread count increases, with the most notable reductions occurring between 1 and 10 threads. Beyond roughly 12-16 threads, the runtime curves begin to level off (or even to rise slightly), reflecting diminishing returns from additional parallelism. This pattern is typical for shared-memory alignment workloads, where the fixed available data bandwidth to the DDR4 memory and parallelization control overhead cause diminishing benefits for scalability at higher thread counts.

RosaSeed consistently achieves the lowest fastq-to-SAM runtime across the full range of application thread counts tested. In single-threaded execution, RosaSeed completes the ERR alignment workload in 699.44 s, while BWA-MEM2 requires 2695.89 s, giving RosaSeed a speedup of approximately 3.85×. RosaSeed also outperforms ERT2, the fastest of the baseline aligners, which takes 1005.14 s under the same single-threaded configuration. This performance advantage carries over to multithreaded execution as well. At 24 threads, RosaSeed finishes in 136.78 s, compared to 209.63 s for BWA-MEM2, 210.50 s for Minimap2, 194.23 s for ERT, and 160.56 s for ERT2, corresponding to speedups of approximately 1.53×, 1.54×, 1.42×, and 1.17×, respectively. Although the runtime gap narrows as more application threads are launched, RosaSeed remains the fastest aligner at every thread count evaluated on the 12-core P620 workstation.

The largest benefits of RosaSeed are observed at lower thread counts, where seed generation costs represent a larger fraction of the total execution time. As the number of application threads grows, memory data bandwidth limitation, fixed cache capacity, cache contention, and other shared-resource bottlenecks become increasingly dominant factors. As a result, the runtime curves of all seven aligners start to converge at higher thread counts.

### 4.6. Energy Consumption

To evaluate energy efficiency, power consumption was measured during the execution of the ERR dataset across application thread counts ranging from 1 to 24. Figure 11 reports the total energy consumed by each aligner in kilowatt-hours (kWh). Power consumption was measured using an inline wall power meter placed between the workstation’s power supply and the AC outlet. The reported energy is the time-integrated power drawn over the full alignment run, including the idle baseline drawn.

**Figure 11.**
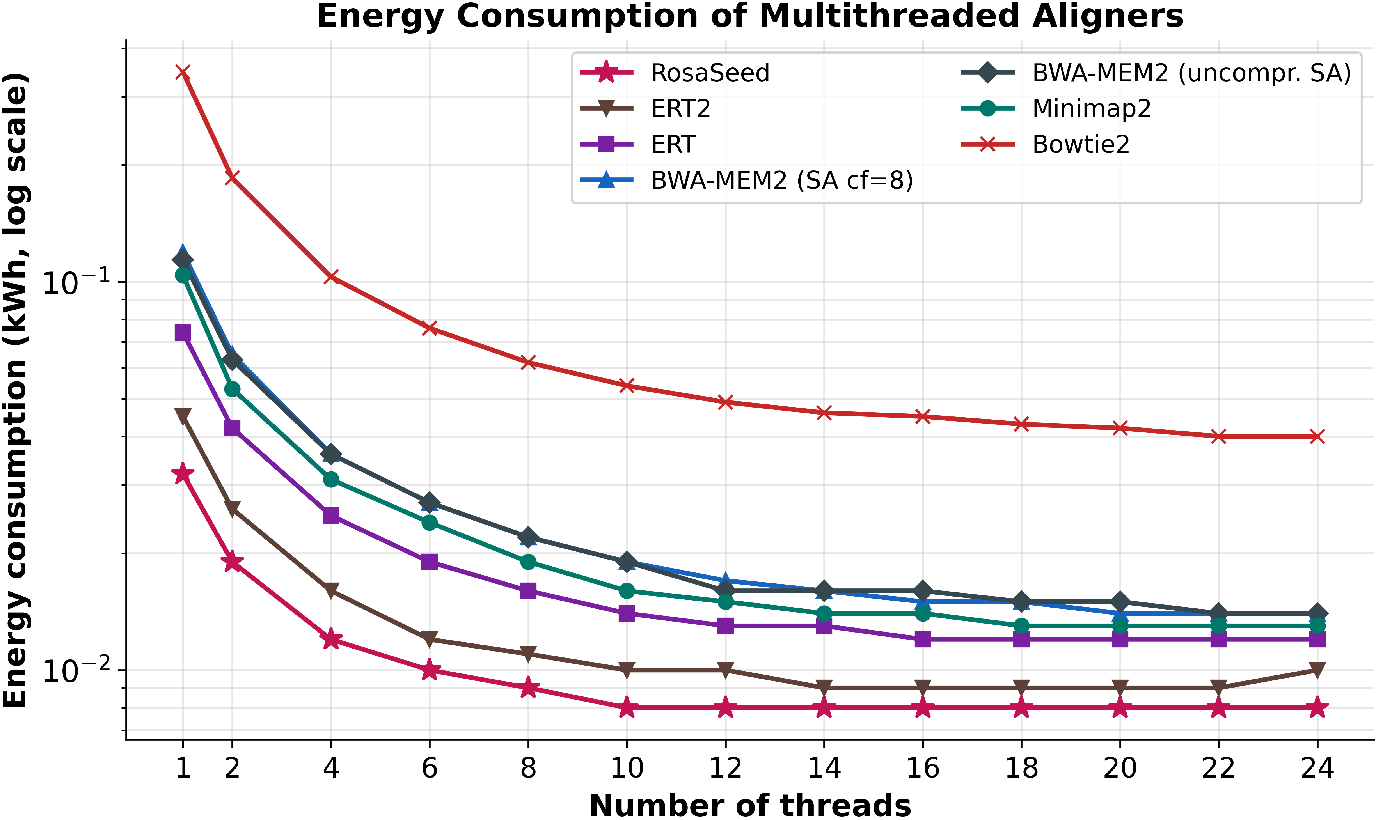
Energy consumption as a function of thread count for the evaluated aligners on the ERR dataset. Each point of the plot is the mean of 12 runs. Energy is reported in kilowatt-hours (kWh) on a logarithmic scale. Lower values indicate greater energy efficiency.

Figure 11 shows that RosaSeed consistently uses the least energy to align the ERR dataset across all tested thread counts. With a single application thread, the workstation running RosaSeed consumes just 0.032 kWh, while the two BWA-MEM2 configurations require 0.114 kWh and 0.119 kWh, respectively. ERT uses 0.074 kWh, ERT2 uses 0.045 kWh, Minimap2 uses 0.104 kWh, and Bowtie2 uses 0.347 kWh. This pattern holds under multithreaded conditions as well. At 24 threads, RosaSeed needs around 0.008 kWh, compared to 0.014 kWh for BWA-MEM2, 0.012 kWh for ERT, 0.010 kWh for ERT2, 0.013 kWh for Minimap2, and 0.040 kWh for Bowtie2. These energy costs closely reflect the runtimes discussed in the previous section. Since RosaSeed significantly reduces the workload involved in seed generation and overall alignment, the processor completes the alignment task in less time, which in turn lowers the total energy needed to align the dataset.

### 4.7. Comparison with minibwa and Strobealign

The experiments in this subsection were conducted on an upgraded workstation (AMD Ryzen Threadripper PRO 5965WX, 24-core Zen 3 microarchitecture, ∼300 GB DDR4) that replaced the Zen 2 platform used for the experiments in Sections 4.1–4.6. The release of minibwa [16] during the preparation of this work allowed us to confirm further that software prefetch-based latency hiding is an effective strategy for accelerating FM index seeding, and that amply supported the coroutine-based scheduling optimization described in Section 3.11. We additionally benchmark against Strobealign [13], a standalone aligner that replaces FM index seeding with fuzzy, subsampled strobemer seeds and that employs its own chaining and extension algorithms. Here we evaluate four configurations: RosaSeed, miniRosaSeed, minibwa and Strobealign, all measured on the ERR dataset, which consists of 20 million 150-bp reads aligned to the T2T-CHM13v2 reference genome.

Table 8 reports per-read runtimes and the same three accuracy metrics for the ERR dataset assuming a single application thread (*t* = 1). With a single thread, miniRosaSeed completes fastq-to-SAM alignment at 13.96 *µ*s/read compared to minibwa’s 29.59 *µ*s/read and Strobealign’s 30.39 *µ*s/read, a 2.12× and 2.18× speedup, respectively, while achieving a standard accuracy of 96.95% versus minibwa’s 94.37% and Strobealign’s 93.85% (margins of 2.58 and 3.10 percentage points). RosaSeed reaches 26.27 *µ*s/read, also faster than both minibwa and Strobealign with a single thread, while achieving 98.52% standard accuracy, the highest of the four configurations and 4.67 percentage points above Strobealign.

**Table 8.** fastq-to-SAM runtime, seed-to-BSW runtime, memory footprint, and alignment accuracy for minibwa, Strobealign, miniRosaSeed, and RosaSeed on the ERR dataset (20M, 150-bp reads, T2T-CHM13v2 reference) for experiments run on a Lenovo P620 AMD 24-core Zen 3 workstation. Runtime metrics are reported in microseconds per read (*µ*s/read) for single-threaded execution (*t* = 1). The *fastq-to-SAM runtime* includes all pipeline stages (index loading, seeding, chaining, extension and output), divided by the number of reads. The *seed-to-BSW* denotes the combined execution time of seed generation, SA lookup, chaining and Banded Smith–Waterman (BSW) extension, divided by the number of reads. Strobealign does not use an FM index or the BSW; for it the corresponding value is the equivalent kernel time (strobemer table lookup, chaining, and Striped Smith–Waterman extension) which is reported as seed-to-BSW runtime.

| Configuration | fastq-to-SAM<br>runtime<br>($\mu\text{s}/\text{read}$) | seed-to-BSW<br>runtime<br>($\mu\text{s}/\text{read}$) | Memory<br>(GB) | Unmapped<br>reads<br>(%) | Std.<br>acc.<br>(%) | Struct.<br>acc.<br>(%) | SCA<br>(%) |
| --- | --- | --- | --- | --- | --- | --- | --- |
| minibwa | 29.59 | 28.18 | 8.30 | 0.391 | 94.37 | 99.94 | 99.84 |
| Strobeatlign | 30.39 | 27.87 | 15.45 | 0.191 | 93.85 | 99.88 | 99.82 |
| miniRosaSeed | 13.96 | 11.20 | 49.61 | 2.094 | 96.95 | 99.96 | 99.83 |
| RosaSeed | 26.27 | 23.07 | 49.61 | 0.523 | 98.52 | 99.97 | 99.87 |

For the seed-to-BSW pipeline, miniRosaSeed requires 11.20 *µ*s/read and RosaSeed 23.07 *µ*s/read, compared to minibwa’s 28.18 *µ*s/read and Strobealign’s 27.87 *µ*s/read. Both RosaSeed configurations are therefore faster than minibwa and Strobealign for the seed-to-BSW kernel runtime as well as the fastq-to-SAM runtime. The difference between fastq-to-SAM and seed-to-BSW kernel time likely reflects pipeline overhead including input/output and index loading, which is larger in RosaSeed due to its 49.61 GB index compared to minibwa’s 8.3 GB and Strobealign’s 15.45 GB indexes.

#### Accuracy

miniRosaSeed produces 4.2× fewer misalignments than minibwa (false positives at ±50 bp: 266,339 versus 1,123,359). Because groundtruth mapping positions are unavailable for the ERR dataset, these counts measure disagreement with BWA-MEM2 rather than confirmed mapping errors. The higher unmapped read fraction of miniRosaSeed (2.09% versus 0.39% for minibwa) reflects a precision– recall trade-off: reads that miniRosaSeed fails to map are likely from alignments to highly repetitive regions of the reference where no seed satisfies the specificity threshold. Note that minibwa aligns these reads but to locations that disagree with BWA-MEM2. In downstream variant-calling workflows, a false-positive alignment can introduce phantom variants into the analysis [37]. Strobealign, despite its fast subsampled seeding, produces the lowest standard accuracy of the four configurations (93.85%). RosaSeed achieves the highest accuracy of the four configurations (standard accuracy 98.52%, structural accuracy 99.97%, sequence-consistent accuracy 99.87%) while leaving only 0.523% of the reads unmapped.

#### Thread Scalability

Figure 12 shows the fastq-to-SAM alignment time as a function of the application thread count (*t*) for two configurations of RosaSeed as well as minibwa and Strobealign. miniRosaSeed is the fastest of the four aligners from *t* = 1 through *t* ≈ 26, where its speed converges with that of minibwa (the two are tied within one standard deviation at *t* ≈ 26). Beyond this point minibwa is marginally faster, reaching a plateau of approximately 32–33 seconds for 20 million reads while miniRosaSeed plateaus at approximately 35–37 seconds. Strobealign, slower than miniRosaSeed throughout in the low-thread range, converges with miniRosaSeed at *t* ≈ 36-38. RosaSeed is faster than minibwa from *t* = 1 through *t* = 6, after which minibwa becomes faster; RosaSeed’s time plateaus at approximately 40–44 seconds.

**Figure 12.**
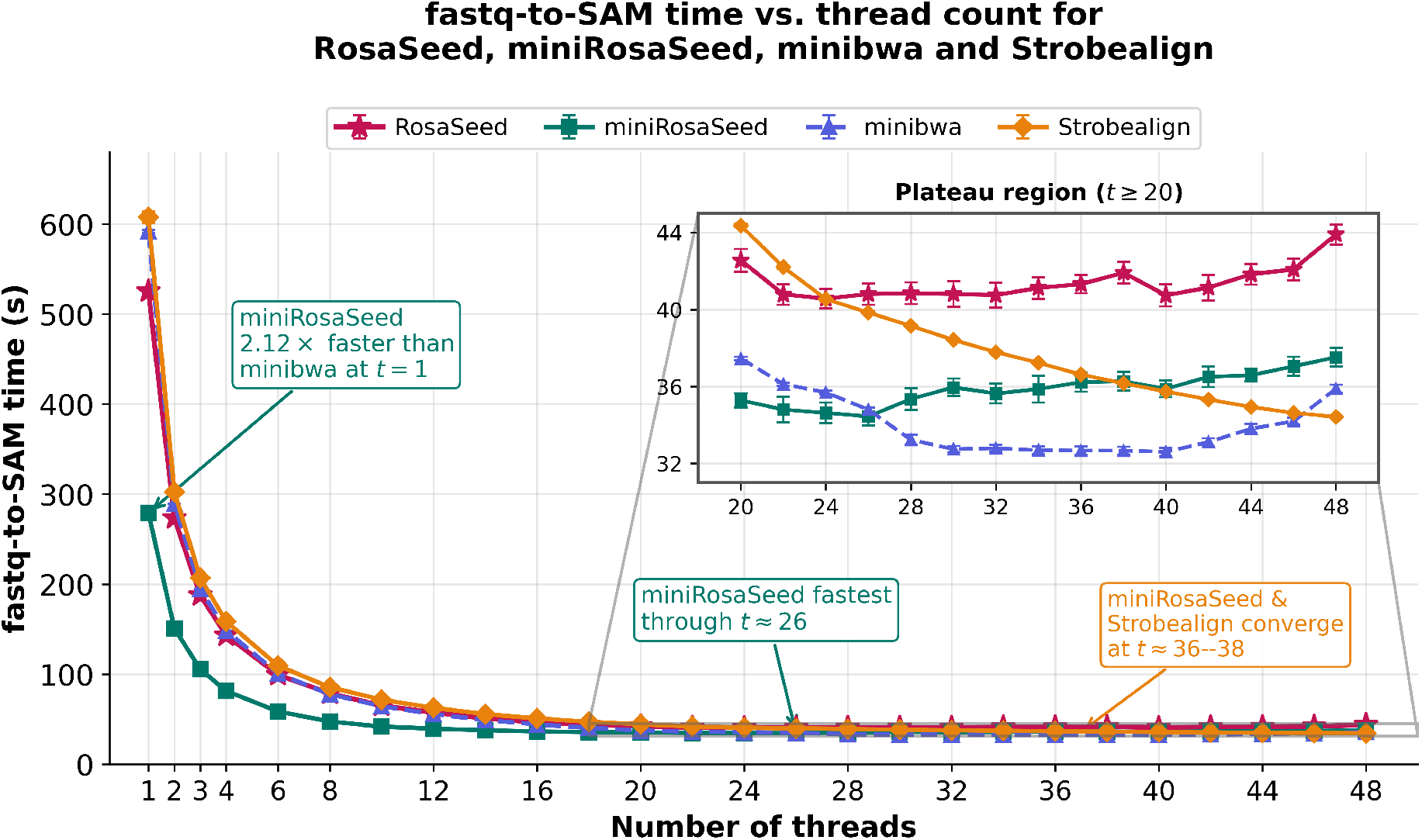
fastq-to-SAM runtime versus application thread count for minibwa, Strobealign, miniRosaSeed, and RosaSeed, measured on the ERR dataset, 150-bp reads aligned to the T2T-CHM13v2 reference genome on a Lenovo P620 AMD Ryzen Threadripper PRO 5965WX workstation (Zen 3, 24 cores). miniRosaSeed is the fastest of the four aligners from *t* = 1 through *t* ≈ 26, with a 2.12*×* speedup over minibwa at a single thread while achieving 2.58 percentage points higher standard accuracy. Beyond *t* ≈ 26 minibwa becomes marginally faster, and miniRosaSeed converges with Strobealign at *t* ≈ 36–38. Recommended RosaSeed is faster than minibwa from *t* = 1 through *t* = 6 and achieves the highest standard accuracy across all thread counts. Error bars denote *±*1 s.d. over 12 runs; minibwa and Strobealign exhibit sub-0.1 s plateau variance, so their bars are smaller than the plotted markers. The inset shows the plateau region (*t* ≥ 20) at an expanded scale.

The plateau observed for all configurations at high thread counts is consistent with a shift from a compute-bound to a resource-bound regime. Two factors likely contribute. First, a portion of the perread pipeline, compressed FASTQ decoding and SAM file output does not parallelise linearly, imposing an effective throughput ceiling independent of seeding speed. Second, the irregular memory-access patterns of large-index alignment place substantial pressure on the fixed available DRAM data bandwidth; as the thread count increases, aggregate bandwidth demand approaches saturation, at which point additional threads yield diminishing returns. This effect is expected to be more pronounced for the RosaSeed configurations, whose 49 GB index far exceeds the cache capacity and is accessed irregularly during seeding, than for minibwa (8.3 GB) or Strobealign (15.45 GB). The lower plateau of both minibwa and the reduced-seed miniRosaSeed configuration is consistent with their smaller downstream workloads and correspondingly lower data bandwidth demand per read.

#### Empirical tuning of coroutine batch size

The coroutine scheduler described in Section 3.11 processes a fixed number of reads concurrently, controlled by the compile-time parameter `RS_BATCH`. This parameter specifies the number of active scheduling slots used to interleave FM index backward-extension steps across reads and therefore determines the degree of memory-latency hiding sought through software prefetching.

To investigate the effect of this parameter on overall alignment performance, we evaluated multiple `RS BATCH` values across alignment thread counts ranging from *t* = 1 to *t* = 48. For each evaluated combination of alignment thread count and `RS BATCH`, the fastqto-SAM runtime was measured over 20 independent runs, and the mean runtime was computed. The results showed a general reduction in the preferred batch size (`RS _BATCH`) as the thread count increased. At low thread counts, larger batches provided greater opportunity to overlap independent FM index memory accesses through software prefetching, thereby improving latency hiding. As additional alignment threads were introduced, the threads themselves generated greater memory-level parallelism, reducing the benefit of large per-thread coroutine batches. Beyond this point, increasing `RS BATCH` further provided little additional latency hiding while potentially increasing contention for shared cache and memory resources. Consequently, progressively smaller `RS BATCH` values were sufficient to achieve the fastest alignment performance at higher thread counts.

Table 9 summarizes the recommended ranges for `RS BATCH` values in RosaSeed. The recommendations group neighboring thread counts rather than reporting the numerically fastest value for each thread count because several adjacent `RS BATCH` values produced statistically similar mean runtimes. For example, for some large thread counts, the difference between the lowest and second-lowest measured mean was substantially smaller than the run-to-run variation. The table should therefore be interpreted as a practical tuning guide rather than as a set of hardware-independent optimal values.

**Table 9.** Recommended ranges for the compile-time RS BATCH parameter on the evaluated AMD Zen 3 workstation. The recommendations represent practical operating ranges rather than hardware-independent optimal settings.

| Alignment threads ($t$) | Recommended `RS_BATCH` values |
| --- | --- |
| 1–4 | 32 |
| 5–12 | 24–28 |
| 13–18 | 12–14 |
| 19–24 | 6–8 |
| 25–30 | 2–6 |
| 31–48 | 1–4 |

The recommendations are specific to the evaluated hardware platform and can be considered as suitable starting points for tuning on other systems. The processor microarchitecture, cache capacity, memory bandwidth, and NUMA configuration will likely influence the exact `RS BATCH` value that produces the shortest runtime. Nevertheless, the overall trend observed across all experiments was consistent: larger `RS BATCH` values provided the best performance at the lower thread counts, whereas progressively smaller `RS BATCH` values became preferable as the thread count was increased.

### 4.8. Downstream Variant Calling Consistency

To evaluate the performance of RosaSeed and miniRosaSeed relative to the established aligners, we benchmarked five alignment pipelines using BWA-MEM2/ERT (both producing the identical SAM output), minibwa, Strobealign, RosaSeed, and miniRosaSeed on the P620 AMD Zen 3 workstation. BWA-MEM2/ERT, minibwa and Strobealign served as reference implementations, whereas RosaSeed and miniRosaSeed are the proposed aligners.

The evaluation was performed using 7,989,074 single-end Illumina HG002 reads (250-bp) aligned against the gapless human reference genome T2T-CHM13v2.0. Variant calling was performed using DeepVariant v1.10.0 [17], and the resulting callsets (the complete set of variants reported by a variant caller) were benchmarked against the GIAB HG002 T2T Q100 v1.1 small-variant truth set [36] using `hap.py` [38] with the `vcfeval` [39] engine. Because this analysis used a subset of the available sequencing data rather than a full-coverage dataset, the results should be interpreted only as an initial benchmark intended to compare the relative behaviour of the five aligners under identical conditions.

#### Alignment statistics

Alignment statistics were obtained using `samtools flagstat` and `samtools stats` and are reported in Table 10 [23]. Strobealign achieved the highest mapping rate (99.91%), followed closely by BWA-MEM2/ERT (99.88%), minibwa (99.86%), and RosaSeed (99.84%). Strobealign did not output supplementary alignment records under the alignment settings used in this study, therefore, its supplementary alignment count is not directly comparable to that of the other aligners. Among the remaining aligners, RosaSeed produced substantially fewer supplementary alignments (49,055) than BWAMEM2/ERT (68,670), representing a 28.6% reduction while maintaining a comparable mapping rate and an alignment error rate (0.004302) nearly identical to that of BWA-MEM2/ERT (0.004300). miniRosaSeed also produced few supplementary alignments (49,154) but exhibited a lower overall mapping rate (98.71%) and the highest alignment error rate (0.004547). The average read length and average base quality were identical across all the aligners because all methods processed the same input reads.

**Table 10.**
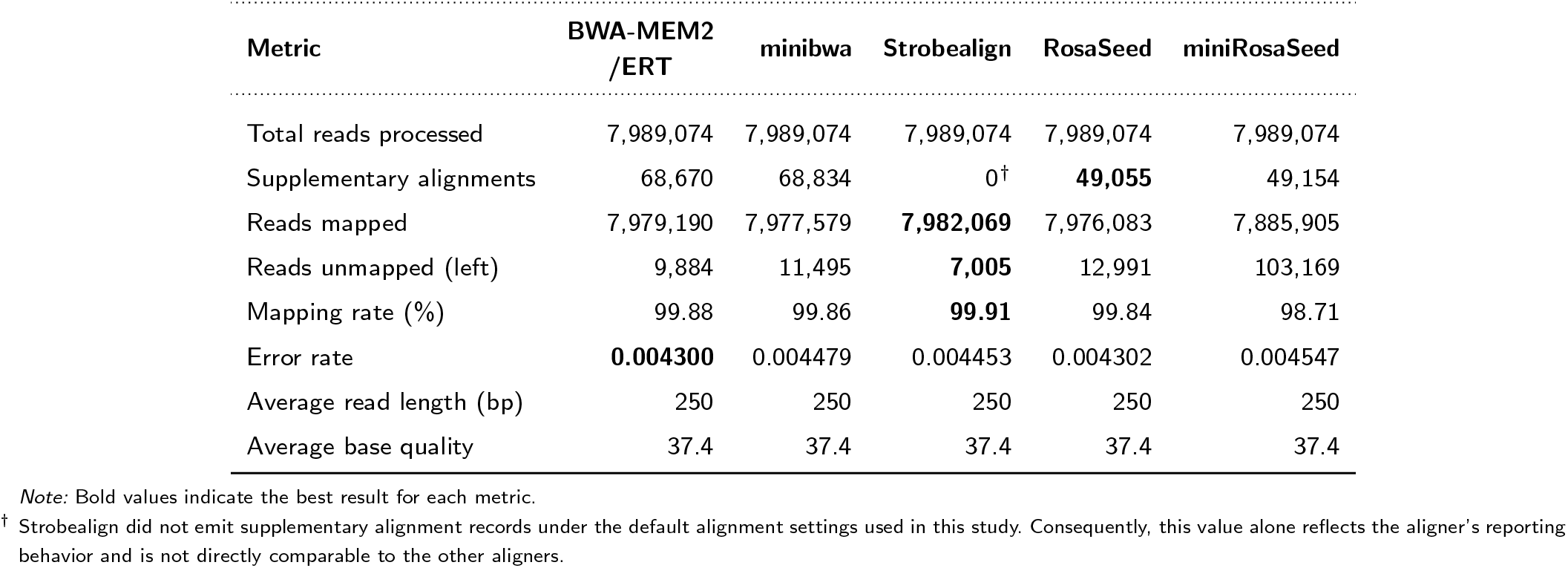
Alignment statistics for BWA-MEM2/ERT, minibwa, Strobealign, RosaSeed and miniRosaSeed on 7,989,074 single-end HG002 Illumina reads (250 bp) aligned to T2T-CHM13v2.0. Supplementary alignment counts and mapping rates were obtained using `samtools flagstat`; all other metrics were obtained using `samtools stats`. The error rate is defined as mismatches divided by mapped bases represented in the CIGAR alignments.

| Metric | BWA-MEM2<br>/ERT | minibwa | Strobealign | RosaSeed | miniRosaSeed |
| --- | --- | --- | --- | --- | --- |
| Total reads processed | 7,989,074 | 7,989,074 | 7,989,074 | 7,989,074 | 7,989,074 |
| Supplementary alignments | 68,670 | 68,834 | 0<sup>†</sup> | **49,055** | 49,154 |
| Reads mapped | 7,979,190 | 7,977,579 | **7,982,069** | 7,976,083 | 7,885,905 |
| Reads unmapped (left) | 9,884 | 11,495 | **7,005** | 12,991 | 103,169 |
| Mapping rate (%) | 99.88 | 99.86 | **99.91** | 99.84 | 98.71 |
| Error rate | **0.004300** | 0.004479 | 0.004453 | 0.004302 | 0.004547 |
| Average read length (bp) | 250 | 250 | 250 | 250 | 250 |
| Average base quality | 37.4 | 37.4 | 37.4 | 37.4 | 37.4 |

Note: Bold values indicate the best result for each metric.
<sup>†</sup> Strobealign did not emit supplementary alignment records under the default alignment settings used in this study. Consequently, this value alone reflects the aligner’s reporting behavior and is not directly comparable to the other aligners.

#### DeepVariant variant discovery

Variants were called independently from each alignment using DeepVariant v1.10.0 [17] under the same workflow. The raw variant counts are shown in Table 11. BWA-MEM2/ERT and minibwa produced nearly identical DeepVariant callsets, differing by only 191 total records. RosaSeed produced 322,897 total records, including 5,062 additional SNPs and 132 additional indels relative to BWAMEM2/ERT. In contrast, Strobealign produced the smallest callset (307,112 total variants), including fewer SNPs and substantially fewer indels (insertions and deletions) than the other aligners, perhaps indicating a more conservative set of variant calls from the downstream DeepVariant analysis. Note that miniRosaSeed produced the largest callset, with 356,058 total records, 38,352 more than BWA-MEM2/ERT, an increase of 12.1% likely driven by 36,832 additional SNPs. This larger miniRosaSeed callset was accompanied by higher numbers of SNP true positives, false positives, and benchmark-unknown calls (variants falling outside the benchmark’s high-confidence regions, which the truth set can neither confirm nor refute) in the subsequent variant benchmarking analysis reported in Table 12.

**Table 11.** Numbers of variant records produced by DeepVariant v1.10.0 from each alignment. Counts were obtained using bcftools stats. Total record counts may not equal the sum of SNP and indel counts because bcftools stats can separately classify a small number of variant records as other variant types.

| Metric | BWA-MEM2<br>/ERT | minibwa | Strobealign | RosaSeed | miniRosaSeed |
| --- | --- | --- | --- | --- | --- |
| Total records | 317,706 | 317,515 | 307,112 | 322,897 | **356,058** |
| SNPs | 282,471 | 282,285 | 275,854 | 287,533 | **319,303** |
| Indels | 35,238 | 35,230 | 31,242 | 35,370 | **36,763** |

Note: Bold values indicate the largest callset in each category.

**Table 12.** SNP benchmarking results against the GIAB HG002 T2T Q100 v1.1 small-variant truth set reference [36] using `hap.py` (`vcfeval` engine). The truth set contains 3,380,403 SNPs. Unknown calls are query variants not classified as true positives or false positives within the benchmark confident regions and are excluded from the precision denominator.

| Metric | BWA-MEM2/ERT | minibwa | Strobealign | RosaSeed | miniRosaSeed |
| --- | --- | --- | --- | --- | --- |
| Truth variants | 3,380,403 | 3,380,403 | 3,380,403 | 3,380,403 | 3,380,403 |
| True positives | 170,467 | 170,656 | 168,298 | 170,878 | **171,090** |
| False negatives | 3,209,936 | 3,209,747 | 3,212,105 | 3,209,525 | **3,209,313** |
| False positives | **66,571** | 66,607 | 65,272 | 66,759 | 67,077 |
| Unknown calls | **14,798** | 15,077 | 16,707 | 18,527 | 31,210 |
| Precision | 0.718896 | 0.719019 | **0.720242** | 0.718806 | 0.718098 |
| Recall | 0.050428 | 0.050484 | 0.049786 | 0.050550 | **0.050612** |
| F1 score | 0.094245 | 0.094344 | 0.093135 | 0.094457 | **0.094560** |
| Fraction not assessed | **0.058811** | 0.059799 | 0.066822 | 0.072388 | 0.115956 |

Note: Bold values indicate the best result for each metric.

#### SNP benchmarking

SNP calls were benchmarked against the GIAB HG002 T2T Q100 v1.1 truth set using `hap.py` [38] with the `vcfeval` [39] engine. The results are shown in Table 12. All five aligners produced broadly similar SNP benchmarking results, although Strobealign exhibited a slightly different performance profile. miniRosaSeed achieved the highest SNP true-positive count (171,090), recall (0.050612), and

F1 score (0.094560), recovering 623 additional true-positive SNPs relative to BWA-MEM2/ERT. RosaSeed achieved the second-highest SNP recall (0.050550) and F1 score (0.094457), whereas Strobealign produced the highest SNP precision (0.720242) but the lowest recall (0.049786), consistent with its smaller and more conservative variant callset.

The overall differences in SNP performance remained small. The F1 scores ranged from 0.093135 (Strobealign) to 0.094560 (miniRosaSeed), indicating that downstream SNP calling was broadly consistent across the evaluated aligners despite modest differences in alignment behaviour and variant discovery.

BWA-MEM2/ERT produced the fewest benchmark-unknown SNP calls (14,798; 5.88% of the SNP callset) whereas miniRosaSeed produced the largest number (31,210; 11.60%). Strobealign generated an intermediate number of benchmark-unknown SNP calls (16,707), reflecting its comparatively smaller DeepVariant SNP callset.

#### Indel benchmarking

Indel results are presented in Table 13. Indel benchmarking results showed similar performance among the BWA-MEM2/ERT, minibwa, RosaSeed, and miniRosaSeed aligners, whereas Strobealign exhibited a more conservative indel calling profile (that is, it produced fewer indel calls overall, reducing false positives at the cost of fewer truepositive calls and lower recall).

**Table 13.**
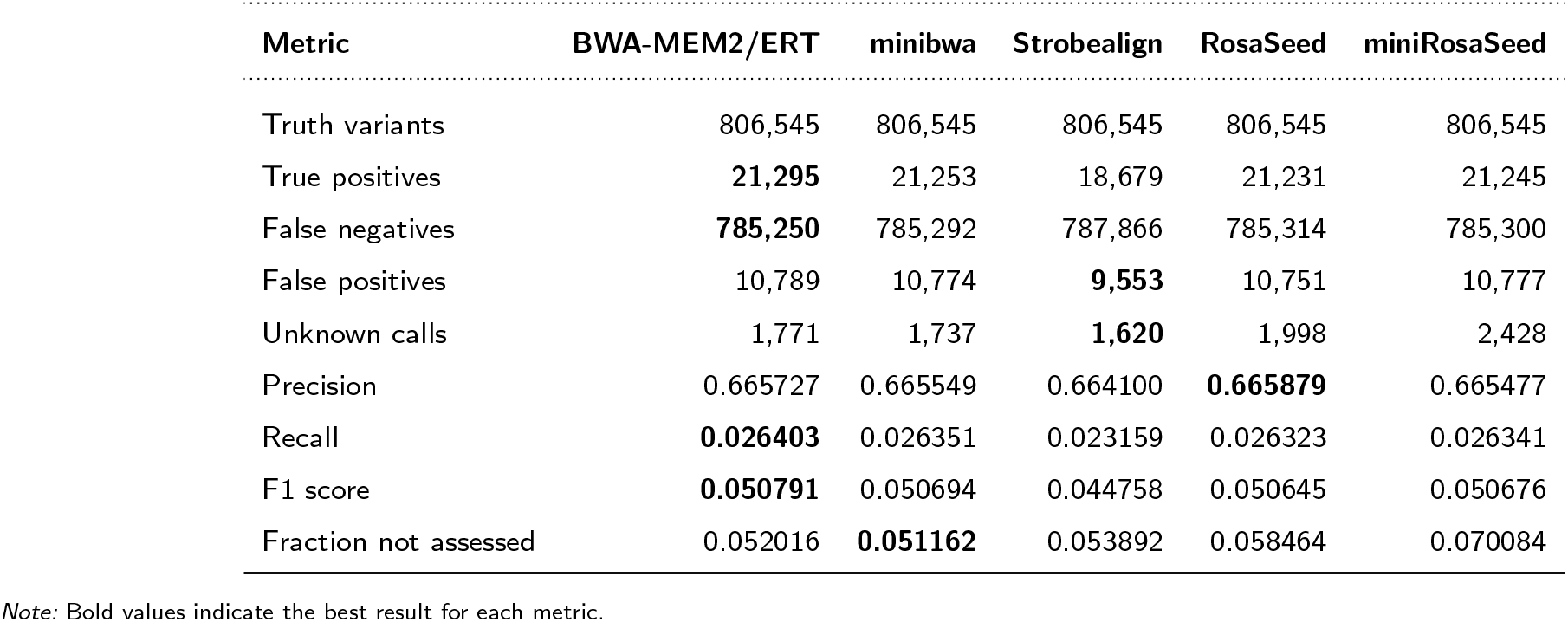
Indel benchmarking results against the GIAB HG002 T2T Q100 v1.1 small-variant truth set [36] using hap.py (vcfeval engine). The truth set contains 806,545 indels. truth set. Strobealign therefore produced fewer false-positive calls but also missed more true variants than the other aligners.

BWA-MEM2/ERT achieved the highest indel true-positive count (21,295), recall (0.026403) and F1 score (0.050791). RosaSeed achieved the highest indel precision (0.665879) while maintaining recall comparable to the BWA-based aligners. In contrast, Strobealign produced the fewest indel false positives (9,553) but also substantially fewer true-positive indels (18,679), resulting in the lowest recall (0.023159) and F1 score (0.044758). These results are consistent with the smaller indel callset generated from the Strobealign alignments.

As with SNPs, miniRosaSeed produced the highest fraction of benchmark-unassessed indel calls (7.01%) compared with 5.20% for BWA-MEM2/ERT.

#### Overall comparison

The five aligners produced broadly comparable downstream variant calling results. The principal differences were associated with their mapping rates, numbers of unmapped and supplementary alignments, mismatch-based alignment error rates, and the size and composition of the DeepVariant callsets generated from those alignments. BWAMEM2/ERT provided the strongest overall baseline, achieving the highest indel recall and F1 score. minibwa closely reproduced the behaviour of BWA-MEM2/ERT, differing by fewer than 200 variant records and by less than 0.0002 in SNP F1 score, demonstrating that it preserves the downstream variant calling characteristics of the original BWA-MEM2 algorithm.

Strobealign followed a different performance profile, achieving the highest mapping rate and the highest SNP precision (0.720242) while producing the smallest DeepVariant callset, particularly for indels. Its higher SNP precision indicates that a slightly larger fraction of its benchmark-assessed SNP calls were true positives. However, its lower SNP recall (0.049786) and indel recall (0.023159) indicate that it recovered a smaller fraction of the variants present in the benchmark

Among the methods evaluated in this work, RosaSeed reduced the number of reported supplementary alignments by 28.6% relative to BWA-MEM2/ERT while maintaining nearly identical mapping statistics and downstream variant-calling performance. Although the RosaSeed-derived callset contained modestly more SNPs than the BWA-based methods, its SNP recall and F1 score remained comparable while achieving the highest indel precision of all five aligners. These results demonstrate that RosaSeed can substantially reduce reported supplementary alignments without compromising downstream variant detection. miniRosaSeed exhibited a different performance profile consistent with its speed-accuracy trade-off: it produced fewer supplementary alignments than the BWA-based methods but had the lowest mapping rate and the highest alignment error rate. DeepVariant generated substantially more SNP calls from the miniRosaSeed alignments, yielding the highest SNP true-positive count, recall, and F1 score. However, the larger callset also contained the highest number of SNP false positives and benchmark-unknown calls, suggesting that the larger callset includes additional variants that were not confirmed within the benchmark confident regions.

Future work will include benchmarking at higher sequencing coverage using paired-end read datasets to determine whether the trends observed in this initial study persist under routine sequencing conditions.

## 5. Discussion

We presented RosaSeed, a fast and memory-efficient short read alignment algorithm that replaces BWA-MEM2’s seeding stage while emitting seeds directly into its downstream chaining and alignment extension pipeline. Using an extensive evaluation on both simulated and real datasets against the T2T-CHM13v2.0 human reference, we showed that both RosaSeed and miniRosaSeed provide significant speedups against the benchmark aligners.

Importantly, these improvements are obtained without modifying BWA-MEM2’s downstream chaining, chain filtering, or Smith– Waterman extension stages, demonstrating that substantial alignment acceleration can be achieved through seed generation alone.

A central observation emerging from the ablation studies is that the Phase A SA-interval cap and Phase C adaptive gap-directed seed generation operate effectively in conjunction. The interval cap suppresses highly abundant seeds that generate substantial downstream processing cost while contributing limited alignment position specificity for chaining. However, pruning away seeds alone will inevitably reduce seed coverage of reads. Phase C compensates for this drawback by directing supplementary seeding effort specifically toward unaligned gap regions, introducing additional pivots to find more seeds only where needed. Table 5 shows that both supplementary strategies are effective, but adaptive gap-directed seeding achieves the best runtime-performance trade-off, reducing the seed-to-BSW time from 38.58 *µ*s/read for fixed-pivot seeding to 29.98 *µ*s/read with essentially identical memory usage. Another interpretation with ERT2 is that, despite eliminating BWA-MEM2’s Phase 2 reseeding pass entirely, ERT2 achieves near-identical accuracy to ERT (99.63% vs. 100.00% standard accuracy on the ERR dataset) while reducing the fastq-to-SAM runtime by 42%. These findings suggest that many seed-level differences are resolved by the downstream alignment stages, resulting in similar final alignments despite differences in the generated seeds.

The coroutine-based prefetch optimization, introduced in response to minibwa [16], yields a further dimension of the speed–accuracy trade-off. miniRosaSeed, which applies more aggressive pre-chain seed suppression and a lower Phase A interval cap (*T* = 50), achieves a 2.12× fastq-to-SAM speedup over minibwa for single-thread execution while maintaining 2.58 percentage points higher standard accuracy and producing 4.2× fewer misalignments, on our 24-core Zen 3 workstation. At thread counts of up to *t* ≈ 26, miniRosaSeed is simultaneously faster and more accurate than minibwa, strictly dominating it on both metrics. At higher thread counts both tools converge to the same input/output-limited plateau (within statistical error), while miniRosaSeed retains its accuracy advantage throughout. The coroutine scheduler itself contributes to a reduction in running times independently of the aggressiveness parameters. These results demonstrate that prefetch-based latency hiding and seed-filtering aggressiveness are orthogonal levers that can be tuned independently to help meet the throughput and accuracy requirements of a given deployment. The same comparison highlights a broader trade-off in subsampling-based seeding: Strobealign, which replaces exact FM index seeds with fuzzy subsampled strobemers, attains the lowest standard accuracy of the evaluated aligners (93.85%) despite competitive speed, whereas miniRosaSeed’s seed filtering heuristic reduces downstream work without abandoning the exact seed foundation, preserving both accuracy and BWA-MEM2 pipeline compatibility.

An initial DeepVariant analysis on 7.9 million single-ended HG002 reads (250-bp) provided additional insight into how alignment differences propagate to downstream variant calls. RosaSeed produced 28.6% fewer supplementary alignments than BWAMEM2/ERT (49,055 versus 68,670) while maintaining a comparable mapping rate (99.84% versus 99.88%) and an essentially identical alignment error rate, suggesting that RosaSeed’s multi-phase seeding produces higher-confidence primary alignments in structurally complex regions where BWA-MEM2 generates split reads (reads whose alignment is divided into two or more separate segments mapped to different reference locations, reported as supplementary alignments). SNP and indel F1 scores were within 0.000315 and 0.000146 of those of BWA-MEM2/ERT, respectively, and RosaSeed achieved the highest indel precision among the five aligners evaluated. Because this experiment used a limited read subset with relatively low genome coverage, these results should be treated as preliminary; they are nonetheless consistent with the broader concordance analysis and confirm that RosaSeed does not introduce systematic errors in downstream variant calling.

RosaSeed’s configurable design accommodates a range of different configurations. RosaSeed-Compact (25.86 GB) suits memory-constrained workstations or shared computing environments. On an AMD Zen 2 workstation platform, the recommended configuration (49.61 GB, 34.97 *µ*s/read) provides the best overall balance of memory, throughput and alignment fidelity for general-purpose sequencing pipelines. RosaSeed-3Base maximizes alignment throughput where memory is plentiful (*>*143 GB). For sensitivity-critical applications, such as rare-variant discovery, clinical reanalysis, or alignment in highly repetitive regions, raising the Phase A SA-interval cap from 2000 to 5000 reduces the unmapped-read fraction from 0.354% to 0.002% on simulated data at a modest increase in runtime cost. The energy measurements further indicate that algorithmic acceleration translates directly into lower computational cost: RosaSeed consumes approximately 3.5× less energy per sample than BWA-MEM2 under single-threaded execution, a consideration of growing operational significance as large-scale genomic initiatives continue expanding in size to include millions of genomes [3, 4].

However, some limitations should be noted. All experiments used the T2T-CHM13v2.0 human reference; the performance of RosaSeed on non-human genomes has not been characterized. Evaluation in this paper was confined to single-ended reads; a paired-ended evaluation on different datasets will be investigated in future work.

Finally, the current implementation is CPU-based, so the reported gains do not exploit hardware acceleration using graphics processing units (GPUs) or field-programmable gate arrays (FPGAs).

Future work will address benchmarking on additional reference genomes including non-human assemblies, and investigation of pangenome graph references, where reduced multi-mapping ambiguity may further enhance the effectiveness of adaptive gap-directed seeding. From a systems perspective, the regularity of RosaSeed’s memory-access pattern and the parallelism of perread jump-table lookups make GPU acceleration a natural next step, offering opportunities to further improve the alignment throughput while maintaining the high accuracy and BWA-MEM2 compatibility. Finally, reads containing ambiguous bases (like `N`) are currently removed during preprocessing as the present implementation supports only the four canonical bases A, C, G and T. Support for reads that contain N’s is planned for future work.

## 6. Conclusions

We presented RosaSeed, a configurable FM index seeding framework for short read alignment that accelerates seed generation through generalized *s*-base references, JT guided initial search, and adaptive supplementary seeding strategies. By replacing only the BWAMEM2 seeding kernel and then feeding candidate seeds directly into its established chaining and Smith–Waterman extension pipeline, RosaSeed achieves 3.85× fastq-to-SAM speedup over BWA-MEM2, 2.48× over ERT and many-fold speedup over other aligners on a single application thread, while requiring approximately 25% less memory than ERT and ERT2, and maintaining close alignment accuracy and downstream variant call concordance comparable to that of BWA-MEM2.

Following the release of minibwa, we introduced a coroutine-based prefetch scheduler that hides FM index cache-miss latency by interleaving extension steps across reads and gap-directed pivots. Building on this optimization, miniRosaSeed, a more aggressive configuration of the RosaSeed, achieves a 2.12× fastq-to-SAM speedup over minibwa for single-thread execution while maintaining 2.58 percentage points higher standard accuracy and producing 4.2× fewer misalignments. miniRosaSeed is the faster of the two through *t* ≈ 26 application threads, beyond which the two converge to roughly the same speed while miniRosaSeed retains its accuracy advantage. At the higher thread counts, the execution times were found to have a large variance, likely due to system-specific activity. When compared with Strobealign, miniRosaSeed is more accurate and 2.18× faster.

The RosaSeed framework exposes multiple runtime-memory operating points through configurable symbol multiplicity (number of bases per symbol), different degrees of SA compression, and supplementary seeding strategies, enabling deployment across memory-constrained workstations and high-throughput production environments alike. These results establish that substantial short read alignment acceleration is achievable through seeding optimization alone, outperforming recent fast aligners, including minibwa and Strobealign, on the accuracy-speed trade-off while preserving alignment quality, memory efficiency, and compatibility with the widely deployed and trusted BWA-MEM2 alignment pipeline.

## Data and Code Availability

The RosaSeed source code is publicly available under the MIT License at https://github.com/smgandhi-18/RosaSeed. The repository also includes the configuration options used in this study to facilitate reproducibility. The T2T-CHM13v2.0 reference genome is publicly available from the Telomereto-Telomere Consortium on NCBI (https://ncbi.nlm.nih.gov/datasets/genome/GCF_009914755.1/), and the real sequencing dataset used for benchmarking is available from the European Nucleotide Archive under accession ERR16657779 (https://www.ebi.ac.uk/ena/browser/view/ERR16657779). Before alignment, all read datasets were preprocessed to remove reads containing ambiguous bases (N); the read counts reported in this study reflect the filtered datasets. The script is available inside the RosaSeed GitHub folder https://github.com/smgandhi-18/RosaSeed/tree/main/preprocessing.

## AI disclosure

AI-assisted language editing tools (Grammarly and Overleaf AI) were used solely to improve grammar, spelling and readability. AI coding assistants were used to assist with drafting utility scripts, routine programming tasks, and improving code readability. All AI-generated code was reviewed, tested, and, where appropriate, modified by the authors before use. All scientific content, algorithm design, software implementation, experimental methodology, data analysis, and conclusions were developed, verified, and approved by the authors, who take full responsibility for the final manuscript and accompanying software.

## 7. Appendix

### 7.1. Notation and configuration parameters

For reference and reproducibility, this appendix consolidates the notation and implementation parameters used throughout the article. Tables A1 and A2 define the symbols used for sequences, FM index operations, seeding phases, and accuracy metrics. Tables A3 and A4 document the compile-time directives and runtime command-line settings used in the reported RosaSeed configuration.

**Table A1.** Notation used throughout this article (part 1): sequences and alphabet, FM index and suffix array, and jump table and SA intervals.

| Symbol | Meaning |
| --- | --- |
| *Sequences and alphabet* |  |
| $F$ | Forward strand of the reference genome (length $L$). |
| $\overline{F}$ | Reverse complement of $F$. |
| $R, \overline{R}$ | Forward read and its reverse complement, sometimes called read query (length $N$). |
| pat | Chosen read strand working sequence used during seeding ($R$ or $\overline{R}$). |
| $N$ | Read length (nt); for example, $N = 150$ for the primary single-end datasets. |
| $\Sigma$ | Nucleotide alphabet $\{A, C, G, T\}$; $\vert \Sigma\vert = 4$. |
| $s$ | Step size (bases per composite symbol) of the $s$-base reference; default $s = 2$. |
| $4^s$ | Size of the $s$-base symbol alphabet (16 symbols for $s = 2$). |
| $F_{\text{even}}, F_{\text{odd}}$ | Even- and odd-phase 2-base symbol sequences of $F$ (for $s = 2$). |
| $\overline{F}_{\text{even}}, \overline{F}_{\text{odd}}$ | Reverse-complement counterparts of the phase-shifted symbol sequences. |
| Ref | Concatenated $s$-base reference used to build the FM index. |
| *FM index and suffix array* |  |
| BWT | Burrows–Wheeler transform of the $s$-base reference. |
| SA[$i$] | Suffix array value at row $i$ (direct, or LF-decompressed if $\text{CF} > 1$). |
| $\sigma$ | A single $s$-base symbol (radix-$4^s$). |
| $\text{Occ}(\sigma, i)$ | Occurrences of symbol $\sigma$ in $\text{BWT}[0, i)$; $\text{Occ}(\sigma, i) = \vert \{j < i : \text{BWT}[j] = \sigma\}\vert $. |
| $C[\sigma]$ | Number of reference positions whose symbol is lexicographically smaller than $\sigma$. |
| $\text{FM}(\sigma, \ell, h)$ | One $s$-base FM backward extension for $\sigma$: $\ell \leftarrow C[\sigma] + \text{Occ}(\sigma, \ell)$, $h \leftarrow C[\sigma] + \text{Occ}(\sigma, h)$, extending the seed by $s$ bases. |
| $\text{CF} = 2^x$ | Suffix array compression factor; only every $\text{CF}$-th SA row is stored. |
| $x$ | Exponent controlling SA compression ($\text{CF} = 2^x$). |
| $p$ | Read position index where noted. |
| *Jump table and SA intervals* |  |
| $JT[q]$ | Jump table entry indexed by the encoded base-level $k$-mer $q$; the table contains $\vert JT\vert = 4^k$ entries. |
| $k$ | Jump table prefix length, with $k \in \{14, 15, 16\}$ nt. |
| $[\ell, h)$ | Suffix array interval; the half-open row range of matching suffixes. |
| $d = h - \ell$ | SA interval width, i.e. the number of genomic hits for the current seed. |
| $d_f, d_r$ | SA interval widths for the forward and reverse-complement read strand at the rightmost $k$-mer. |
| $\text{RefWalk}(p, p_r)$ | Left extend a seed from read position $p - k$ and reference position $p_r$; returns additional matched bases. |
| seed_len | Current seed length (nt) during extension. |
| $i$ | Generic position index. |

**Table A2.** Notation used throughout this article (part 2): seeding parameters (Phases A–C) and accuracy metrics.

| Symbol | Meaning |
| --- | --- |
| *Seeding parameters (Phases A–C)* |  |
| $T$ | Phase A SA-interval cap; seeds with $d > T$ are suppressed. $T = 0$ disables the cap. |
| $\ell_A = 19$ nt | Minimum seed length emitted in Phase A. |
| $\ell_{BC} = 20$ nt | Minimum seed length emitted in Phase B and Phase C ($\ell_{BC} = \ell_A + 1$). |
| $\delta = 20$ | Specificity threshold: maximum SA interval width for Phase B/C seed emission. |
| $s_{\text{GF}}$ | Gap-directed pivot step size: 2 if the gap $\leq \ell_{BC}$ (20 nt), otherwise 4. |
| $\Gamma = 5$ nt | Minimum gap size that triggers Phase C gap-directed seeding. |
| $G = [g_{\text{start}}, g_{\text{end}}]$ | An unseeded gap interval on the read; $g = g_{\text{end}} - g_{\text{start}} + 1$ is its length. |
| $S_A$ | Phase A output seed list. |
| $S_{AB}, S_{AC}$ | Combined seed output of Phase A and Phase B, Combined seed output of Phase A and Phase C. |
| *Accuracy metrics* |  |
| $a^*, a$ | Ground truth or reference alignment and aligner-reported alignment for a read, respectively. |
| $\text{start}(a^*), \text{start}(a)$ | Leftmost start positions of the truth and reported alignments. |
| $I(r), D(r), X(r)$ | Number of insertions, deletions, and mismatches in alignment $r$. |
| $E(r) = I(r) + D(r) + X(r)$ | Total edit count of alignment $r$. |
| $\text{NM}(a)$ | Edit distance (SAM NM tag) of the reported alignment $a$ to the reference. |

**Table A3.**
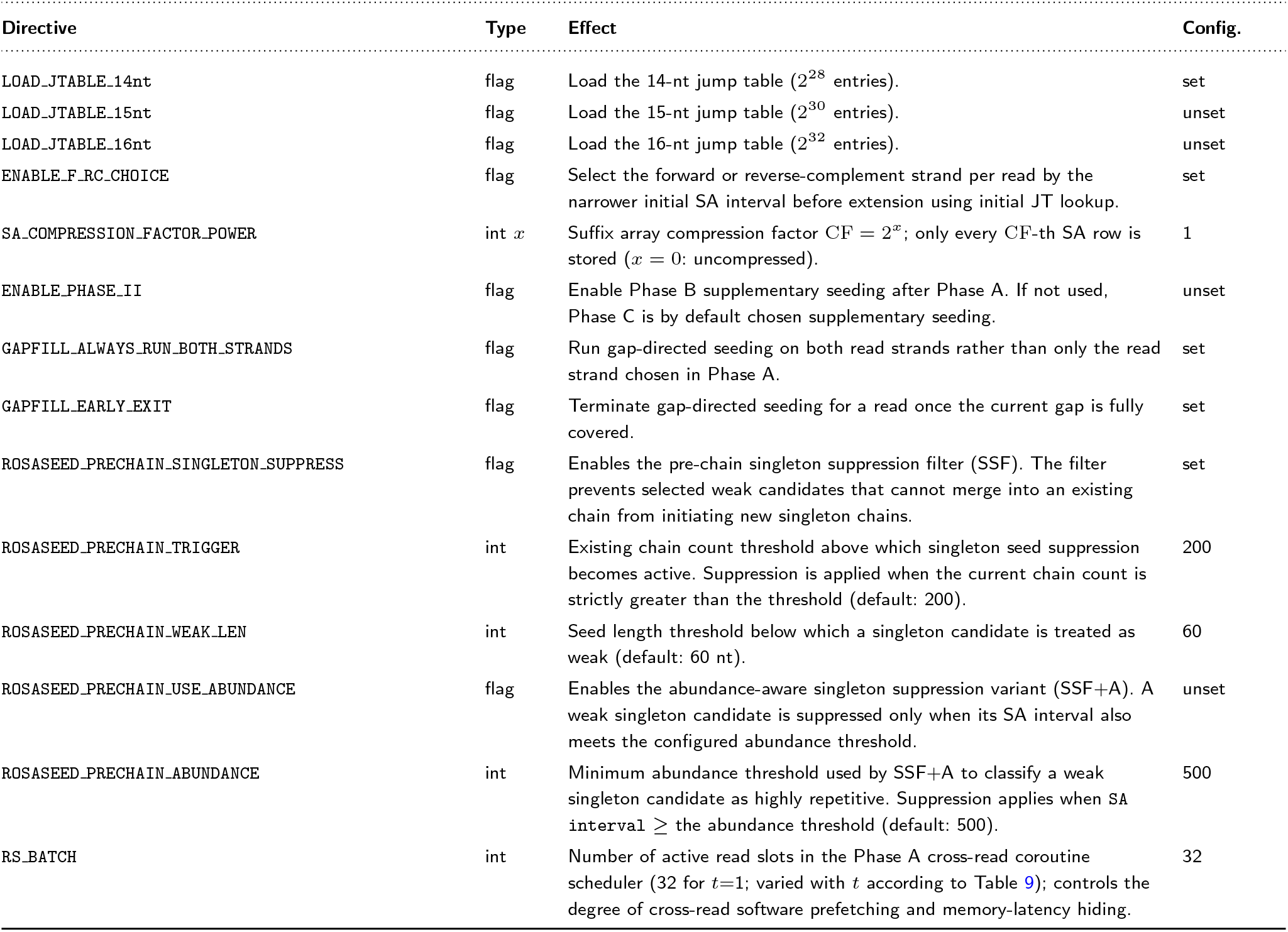
Compile-time preprocessor directives controlling the RosaSeed build. Directives are supplied through CPPFLAGS EXTRA at make time (e.g. -DLOAD JTABLE 14nt). The final column lists the value used for the *RosaSeed* configuration reported in this article.

| Directive | Type | Effect | Config. |
| --- | --- | --- | --- |
| LOAD_JTABLE_14nt | flag | Load the 14-nt jump table ($2^{28}$ entries). | set |
| LOAD_JTABLE_15nt | flag | Load the 15-nt jump table ($2^{30}$ entries). | unset |
| LOAD_JTABLE_16nt | flag | Load the 16-nt jump table ($2^{32}$ entries). | unset |
| ENABLE_F_RC_CHOICE | flag | Select the forward or reverse-complement strand per read by the narrower initial SA interval before extension using initial JT lookup. | set |
| SA_COMPRESSION_FACTOR_POWER | int $x$ | Suffix array compression factor $CF = 2^x$; only every $CF$-th SA row is stored ($x = 0$: uncompressed). | 1 |
| ENABLE_PHASE_II | flag | Enable Phase B supplementary seeding after Phase A. If not used, Phase C is by default chosen supplementary seeding. | unset |
| GAPFILL_ALWAYS_RUN_BOTH_STRANDS | flag | Run gap-directed seeding on both read strands rather than only the read strand chosen in Phase A. | set |
| GAPFILL_EARLY_EXIT | flag | Terminate gap-directed seeding for a read once the current gap is fully covered. | set |
| ROSASEED_PRECHAIN_SINGLETON_SUPPRESS | flag | Enables the pre-chain singleton suppression filter (SSF). The filter prevents selected weak candidates that cannot merge into an existing chain from initiating new singleton chains. | set |
| ROSASEED_PRECHAIN_TRIGGER | int | Existing chain count threshold above which singleton seed suppression becomes active. Suppression is applied when the current chain count is strictly greater than the threshold (default: 200). | 200 |
| ROSASEED_PRECHAIN_WEAK_LEN | int | Seed length threshold below which a singleton candidate is treated as weak (default: 60 nt). | 60 |
| ROSASEED_PRECHAIN_USE_ABUNDANCE | flag | Enables the abundance-aware singleton suppression variant (SSF+A). A weak singleton candidate is suppressed only when its SA interval also meets the configured abundance threshold. | unset |
| ROSASEED_PRECHAIN_ABUNDANCE | int | Minimum abundance threshold used by SSF+A to classify a weak singleton candidate as highly repetitive. Suppression applies when SA interval ≥ the abundance threshold (default: 500). | 500 |
| RS_BATCH | int | Number of active read slots in the Phase A cross-read coroutine scheduler (32 for $t=1$; varied with $t$ according to Table 9); controls the degree of cross-read software prefetching and memory-latency hiding. | 32 |

**Table A4.** Runtime command-line flags for the RosaSeed-integrated bwa-mem2 mem command. The *Default* column is the built-in value used when the flag is omitted; the *Config*. column is the value used for the recommended RosaSeed configuration reported in this article.

| Flag | Arg | Meaning | Default | Config. |
| --- | --- | --- | --- | --- |
| --rs-index | STR | Path to the pre-built RosaSeed index directory (required in RosaSeed mode). | — | *index dir* |
| --rs-cap | INT | Phase A SA-interval cap $T$; seeds whose interval width exceeds $T$ are pruned away. | 2000 | 2000 |
| --rs-gap | INT | Minimum gap size $\Gamma$ (bp) that triggers Phase C adaptive gap-directed recovery. | 5 | 5 |
| --rs-pivots-b | INT | Number of fixed pivots placed in Phase B supplementary seeding. | 3 | 3 |
| --rs-gap-step | INT | Pivot step size (bp) for gap-directed left-extension in Phase C. | 4 | 4 |
| *Inherited BWA-MEM2 flags used in the RosaSeed runs* |  |  |  |  |
| -t | INT | Number of worker threads. | 1 | 1 |
| -k | INT | Minimum seed length passed to $\ell_A$ in RosaSeed. $\ell_{BC} = \ell_A + 1$. | 19 | 19 |
| -y | INT | SA interval width threshold for supplementary seeding passed to $\delta$ in RosaSeed. | 20 | 20 |

## Footnotes

1 Coverage gives the average number of times each base in the genome is sequenced; at 5*×*, each position is covered by five reads on average. For the *∼*3.1 Gbp T2T-CHM13v2.0 assembly, this corresponds to 5 *×* Gbp*/*150 bp ≈ 103 million reads.

